# CXCL16 mediates nociception and inflammation in murine post-traumatic osteoarthritis

**DOI:** 10.1101/2025.04.10.647297

**Authors:** Lindsey Lammlin, Stephen Redding, Alexander J. Knights, Aanya Mohan, Michael D. Newton, Rachel F. Bergman, Isabelle J. Smith, Scarlet C. Howser, Sofia Gonzalez-Nolde, Phillip M. Rzeczycki, Natalie S. Adamczyk, Rachel E. Miller, Tristan Maerz

**Affiliations:** ETH Zürich, Zürich, Switzerland; Department of Orthopaedic Surgery, University of Michigan; Ann Arbor, Michigan, United States; Washington University St. Louis, St. Louis, Michigan; Division of Rheumatology, Rush University Medical Center; Chicago, Illinois, United States; Department of Biomedical Engineering, University of Michigan; Ann Arbor, Michigan, United States; Division of Rheumatology, Department of Internal Medicine, University of Michigan; Ann Arbor, Michigan, United States

**Author notes:** Corresponding Author Info Tristan Maerz, PhD ETH Zürich, Zürich, Switzerland. Gloriastrasse 39, 8092 Zürich, Switzerland.

## Abstract

This study investigates the role of the chemokine CXCL16 and its receptor, CXCR6, in post-traumatic osteoarthritis (PTOA) and joint nociception, highlighting the potential of targeting the CXCL16-CXCR6 axis for therapeutically managing joint inflammation and pain. Following joint injury in mice, the CXCL16-CXCR6 signaling axis is activated in synovium, driven by synovial fibroblasts and macrophages. Human OA synovium also exhibited increased CXCL16 and CXCR6 gene expression. CXCL16 stimulated a pro-inflammatory response in fibroblasts and macrophages, contrasting with an anti-inflammatory response observed in mesenchymal progenitor cells. In mice, repeated intra-articular CXCL16 injections induced histological synovitis and sex-dependent activation of inflammatory and fibrotic transcriptional programs in synovium. Repeated CXCL16 joint injections also induced knee hyperalgesia, which was mitigated by co-administration of the CXCR6 antagonist, ML339. A single intra-articular injection of CXCL16 induced acute knee hyperalgesia as early as 30 minutes post-injection, which was completely abrogated by ML339 co-treatment, suggesting direct CXCL16 binding to nociceptor-expressed CXCR6. In a murine PTOA model, systemic CXCR6 antagonism with ML339 alleviated knee hyperalgesia and altered circulating immune cell profiles. Direct stimulation of mouse dorsal root ganglion-derived nociceptive neurons with CXCL16 induced rapid calcium signaling, which was abolished by co-treatment with ML339. These findings establish CXCL16 as a regulator of joint inflammation and identifies the CXCL16-CXCR6 binding mechanism as key in mediating pain-related behaviors and nociceptor activation, offering a therapeutic target for PTOA-related inflammation and pain management.

**One Sentence Summary:** CXCL16 regulates synovial inflammation and mediates joint nociception via CXCR6, highlighting its potential as a therapeutic target for post-traumatic osteoarthritis.

## INTRODUCTION

The role of synovitis in promoting multiple pathological manifestations of osteoarthritis (OA), including pain and catabolic articular destruction, is now well-recognized(*1*, *2*). In idiopathic OA, synovitis manifests as chronic low-grade tissue inflammation with intermittent flare-ups, and recent literature has demonstrated the existence of different inflammatory endotypes in both late-stage OA and rheumatoid arthritis (RA)(*3–5*). In the setting of post-traumatic osteoarthritis (PTOA), joint injury results in a high degree of acute synovial inflammation, which transitions to a chronic, low-grade nature akin to that observed in idiopathic OA.

A highly coordinated and tightly regulated set of factors orchestrate the local and systemic responses to tissue trauma. However, the immunological mechanisms underpinning joint injury-induced synovitis and its downstream consequences remain poorly described. Among the factors strongly induced following injury are the families of chemokines, secreted by tissue-resident stromal, endothelial, and immune cells. While some chemokines such as CCL2/MCP-1(*6–10*) have been shown to be involved in OA/PTOA inflammation, the individual functions of most chemokines remain unclear in the context of joint injury and PTOA development. For instance, while CCL5/RANTES was shown to activate IL6 production in human synovial fibroblasts(*11*), indicating a potential deleterious pro-inflammatory effect, its genetic ablation did not alter disease severity in a surgical mouse OA model(*12*). Here, we identified and functionally characterized the chemokine CXCL16 as a synovial-derived factor selectively upregulated and secreted following joint injury. CXCL16 is expressed as a transmembrane protein, and it is cleaved from the cell surface by a disintegrin and metalloproteinase 10 (ADAM10) to act as a soluble ligand(*13*). As a transmembrane protein, CXCL16 is thought to act as an adhesion molecule for leukocytes(*14*) and can also act as a receptor for the soluble ligand form of CXCL16(*15*, *16*). CXCL16 also binds CXCR6, a G-protein coupled receptor and the only classical receptor described to interact with CXCL16(*17*, *18*). The CXCL16-CXCR6 axis has been associated with *in vivo* chemotaxis of macrophages, MSCs, and T cells in lung, liver, cardiac, renal, skin, and rheumatoid arthritic joint tissues(*19–24*). In the context of RA, CXCL16 has been shown to recruit mononuclear cells to synovial tissue(*25*) and promote FLS proliferation(*26*). CXCL16 has also been shown to be involved in the recruitment of endothelial progenitor cells, contributing to pathological angiogenesis observed in RA(*27*). Ablation of CXCR6 ameliorated K/BxN serum-induced arthritis by mitigating inflammatory cell infiltration, and fewer T cells and macrophages were found in joints of K/BxN-challenged CXCR6^-/-^ mice(*27*). CXCL16 monoclonal antibody treatment was also shown to be effective in mitigating collagen induced arthritis in mice(*28*). Despite this evidence from inflammatory and autoimmune-mediated arthritis, no prior study has described the role of the CXCL16-CXCR6 axis following joint injury and during PTOA progression.

Chemokines are also highly relevant in the context of pain, and pain mechanisms in OA/PTOA remain poorly described. Chemokines can either directly affect nociception as agonists of nerve fiber-bound receptors, serving to sensitize nociceptive transmission underpinning pain, or indirectly by inducing the upregulation of cytokines and neurotrophic mediators that interact with neurons(*29*, *30*). Very few chemokines have been characterized in their involvement in joint nociception and OA/PTOA-related pain specifically. In a mouse model of gouty arthritis, the CXCL5-CXCR2 axis was shown to have a role in pain and neurogenic inflammation as mice with neuronal specific CXCR2 depletion exhibited less severe pain related-behaviors and reduced production of calcitonin gene-related peptide and substance P(*31*). Prior work has demonstrated that CCL2 is a key mediator of joint nociception by interacting with nociceptor-bound CCR2(*6*). Furthermore, CCL2^-/-^ and CCR2^-/-^ mice exhibit delayed onset of pain-related behavior in a surgical PTOA model(*7*, *8*). The CXCL16-CXCR6 axis, however, has yet to be described in the context of joint nociception.

The objective of this study was to assess the role of CXCL16 in joint injury-induced, PTOA-associated inflammation and nociception. We demonstrate that the CXCL16-CXCR6 axis is activated following traumatic joint injury, that CXCL16 induces pro-inflammatory signaling in synovial fibroblasts and macrophages, and that CXCL16 is sufficient by itself to induce joint pain and synovitis in mice *in vivo*. Using *in vitro* Ca^2+^ measurements in neuronal cells derived from nociceptor-specific GCaMP6s calcium sensor mice and using *in vivo* acute hyperalgesia experiments, we show that CXCL16 directly agonizes nociceptors and induces knee hyperalgesia. Finally, employing a clinically-relevant pharmacological CXCR6 antagonist, we demonstrate that CXCR6 inhibition mitigated the pro-algesic effects of CXCL16, abrogated CXCL16-induced Ca^2+^ signaling by nociceptors, and ameliorated joint injury-induced knee hyperalgesia.

## RESULTS

### Traumatic joint injury activates the CXCL16-CXCR6 axis in synovium

To understand the temporal expression patterns of synovial-derived chemokines and their receptors following joint injury, we analyzed our bulk RNA-sequencing (RNA-seq) data of whole mouse synovium (inclusive of Hoffa’s fat pad)(*32*). We observed marked injury-induced perturbation in members of the CCL and CXCL families (**Fig. 1A**), with corresponding changes in the expression pattern of their respective receptors (**Fig. S1**). We next screened this list for chemokines with zero or near-zero expression in Sham joints, further focusing on chemokines with no prior PTOA-related studies. We identified CXCL16, which was upregulated 4.22- and 2.97-fold at 7d and 28d post-ACLR, respectively, with markedly lower *P_adj_* values compared to all other chemokines (**Fig. 1A**). qPCR corroborated the upregulation of *Cxcl16* at additional timepoints post-ACLR in the synovium and meniscus (**Fig. S2A-B**). CXCL16 is expressed as a transmembrane protein, in which state it can function as a scavenger receptor for lipoproteins and bacteria(*33*) in addition to acting as a receptor for the soluble form of CXCL16(*15*, *16*).

**Fig. 1.**
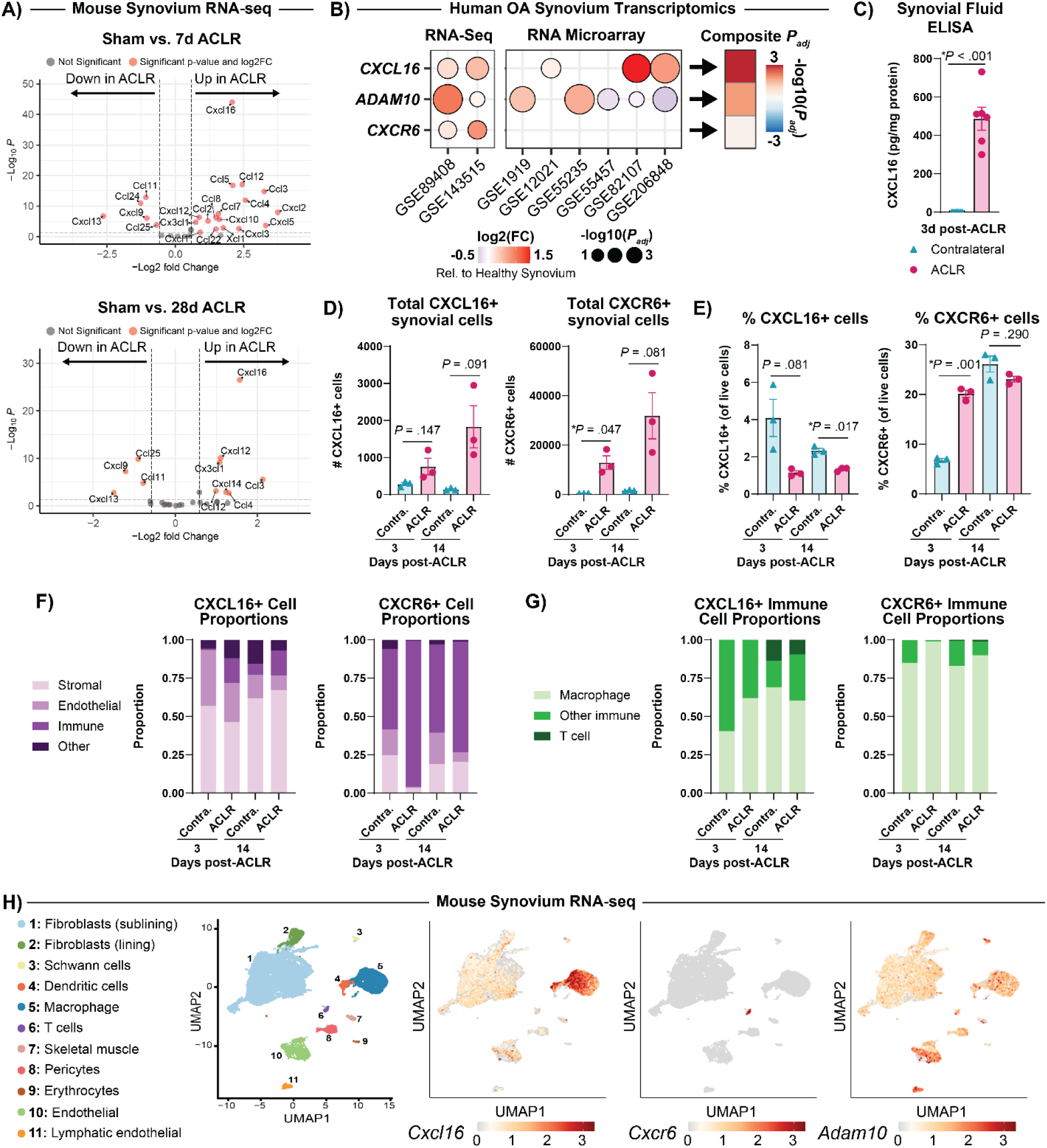
Joint injury activates the CXCL16-CXCR6 axis in synovium. (**A**) Volcano plots of mouse synovial bulk RNAseq data(*32*) comparing chemokine ligand expression in Sham vs 7d ACLR and Sham vs 28d ACLR. Genes with red datapoints were differentially expressed at *P_adj_* < 0.05 and |log_2_FC| > 0.585. (**B**) Bubble plot from a meta-analysis of human OA synovial transcriptomics datasets showing expression of *CXCL16*, *CXCR6*, and *ADAM10* relative to healthy synovium. (**C**) CXCL16 ELISA of synovial fluid in contralateral and injured joints 3d post-ACLR (n=6 per group). Total number of (**D**) and percent positive (**E**) CXCL16+ and CXCR6+ cells in contralateral and injured synovium at 3d and 14d post-ACLR (n=3 per group. 1 sample = 1 male + 1 female synovia pooled). (**F**) Proportions plots showing the cell type breakdown of CXCL16+ and CXCR6+ cells. (**G**) Proportions plots further breaking down the CXCL16+ and CXCR6+ immune cell types. (**H**) Feature plots of mouse synovial single cell RNAseq data(*35*) showing *Cxcl16, Cxcr6,* and *Adam10* gene expression distribution across all cell types. All bars show mean ± SEM.

Upon cleavage by ADAM10, CXCL16 is shed from the membrane and acts as a soluble chemokine, shown to mediate leukocyte chemotaxis(*14*) as well as activate the phosphatidylinositol-4,5-bisphosphate 3-kinase (PI3K)-Akt/PKB and extracellular signal-regulated kinase mitogen-activated protein kinase (ERK/MAPK) pathways, and mobilize Ca^2+^ via the G protein-coupled receptor CXCR6(*34*). Mining of our previously published bulk RNA-seq data of whole synovium also showed that *Cxcr6* was upregulated 3.30-fold at 7d and 2.87-fold at 28d post-ACLR (**Fig. S1A**). By qPCR we also observed upregulation of *Adam10* in the synovium, but not the meniscus following joint injury (**Fig. S2A-B**), demonstrating upregulation of the entire CXCL16-CXCR6-ADAM10 axis following joint injury.

In a meta-analysis of transcriptomic data of human OA synovium, we also observed marked alterations in the expression patterns of CCL and CXCL chemokines and their receptors, relative to healthy synovium (**Fig. S1B-C**). *CXCL16*, *CXCR6*, and *ADAM10* were significantly upregulated in OA synovium compared to healthy synovium (**Fig. 1B**). Notably, CXCL16 had the lowest, i.e. most significant composite *P_adj_* value compared to all other CCL and CXCL chemokines (**Fig. S1B**), demonstrating that CXCL16 upregulation is also observed in human OA and not confined to the acute post-injury period. Uninjured mouse joints and healthy human OA synovium exhibited very low CXCL16 expression. Concordantly, while CXCL16 protein was undetectable in the synovial fluid of uninjured joints, the synovial fluid of injured ACLR joints was enriched in CXCL16 protein (**Fig. 1C**).

To identify the synovial cells expressing CXCL16 (intracellular) and CXCR6 (surface) proteins, we performed flow cytometry of ACLR synovium compared to healthy contralateral synovium at both early (3d) and intermediate (14d) timepoints following injury. We observed large but statistically non-significant increases in the total number of CXCL16+ and CXCR6+ cells in ACLR synovia compared to uninjured synovia (**Fig. 1D**), but only CXCR6+ cells increased in their proportion to all live cells at 3d post-ACLR (**Fig. 1E**). This is likely due to the widely reported dramatic increase in total synovial cells after joint injury, corroborated by our results showing all major synovial cell types elevated at 3d and 14d post-ACLR compared to contralateral tissue (**Fig. S2C-G**). Intracellular CXCL16 was found to be expressed most strongly by CD31(-) CD45(-) stromal cells, i.e. synovial fibroblasts, as well as by CD31(+) CD45(-) endothelial cells, and by CD45(+) immune cells (**Fig. 1F**). Within CD45(+) immune cells, we observed CXCL16 expression in CD11b(+) CD64(+) macrophages, CD11b(-) CD3(+) T cells, and other immune subsets (**Fig. 1G**). CXCR6 was predominantly expressed by immune cells with lower levels of expression in stromal and endothelial cells (**Fig. 1F**). The CXCR6+ immune cells were comprised of ∼80-90% macrophages (**Fig. 1G**). Mining of our previously published murine synovial single-cell RNA sequencing data(*35*) shows that *Cxcl16* transcript is expressed by fibroblasts and most highly in macrophage and dendritic cells, whereas *Cxcr6* transcripts are confined to T-cells, and *Adam10* is expressed ubiquitously across synovial cell populations (**Fig. 1H**).

Taken together, these findings demonstrate that CXCL16 is a disease-specific secreted factor, expressed by synovial fibroblasts and macrophages, and enriched in the synovium of both human OA patients and injured mouse joints.

### IL1β and TNFα induce CXCL16 expression in joint relevant cell types

Given the observation of injury-induced upregulation of CXCL16 by multiple synovial cell types, we sought to identify upstream activators of CXCL16 expression (**Fig. 2A**). We found that IL-1β and TNFα, two key cytokines strongly upregulated in synovium following ACLR(*32*, *36*), activated *Cxcl16* and *Cxcr6* transcript upregulation in both primary murine FLS and bone marrow-derived macrophages (**Fig. 2B-C**). IL-1β and TNFα also caused upregulation of *Adam10* in FLS (**Fig. 2B**), but downregulation of *Adam10* in BMDM (**Fig. 2C**). We confirmed the increase of secreted CXCL16 protein in the conditioned medium of IL-1β- and TNFα-treated FLS, compared to vehicle-treated cells (**Fig. 2B**). We also found IL-1β and TNFα to cause upregulation of *Cxcr6* in an immortalized chondrogenic cell line, ATDC5s - a culture model of articular chondrocytes, whereas these cytokines induced downregulation of *Adam10* while *Cxcl16* remained unchanged (**Fig. 2D**). These results demonstrate that the CXCL16-CXCR6 axis is activated by IL1β and TNFα.

**Fig. 2.**
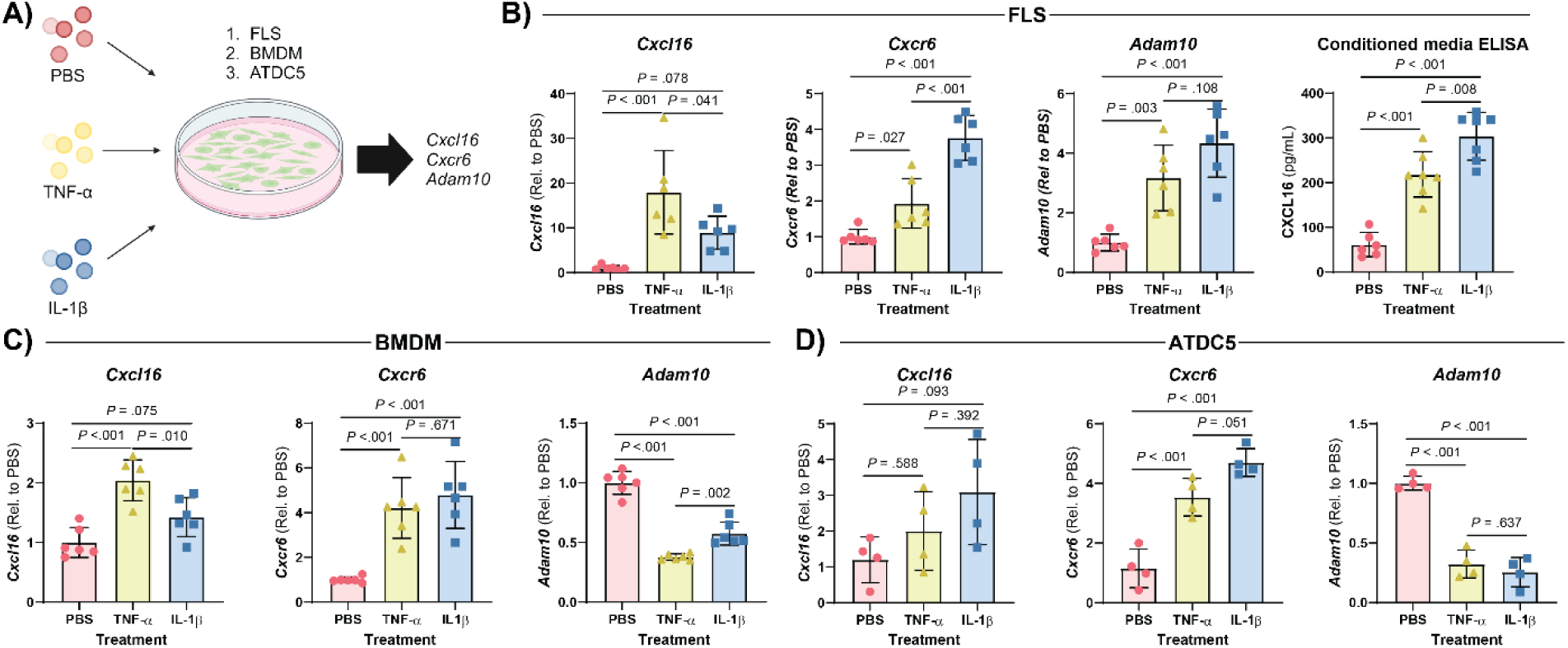
Upstream mediators of the CXCL16 signaling axis *in vitro.* (**A**) Fibroblast like synoviocytes (FLS), bone marrow-derived macrophages (BMDM), and immortalized chondrogenic cells (ATDC5s) were treated with pro-inflammatory cytokines and perturbation of the CXCL16 signaling axis was evaluated via qPCR. (**B**) Gene expression of *Cxcl16*, *Cxcr6*, and *Adam10* by qPCR in FLS (n=6 per group) and a CXCL16 ELISA of FLS conditioned media (n=6-7 per group). (**C**) Gene expression of *Cxcl16*, *Cxcr6*, and *Adam10* by qPCR in BMDMs (n=6 per group). (**D**) Gene expression of *Cxcl16*, *Cxcr6*, and *Adam10* by qPCR in ATDC5s. (n=4 per group). All bars show mean ± SEM.

### CXCL16 induces cell type-dependent pro- or anti-inflammatory gene expression

We next wanted to assess the potential functional role of CXCL16 in joint-relevant cell types. We treated primary murine synovial fibroblasts, bone marrow-derived macrophages, and bone marrow-derived mesenchymal progenitor cells (BMPCs) with increasing doses of recombinant CXCL16 protein. We found CXCL16 to induce a multi-factorial pro-inflammatory response in FLS, indicated by both an increase in pro-inflammatory gene expression (*Il1b, Il6*) and a decrease in anti-inflammatory gene expression (*Il10*) (**Fig. 3A**). BMDMs treated with CXCL16 similarly had a similar double-edge pro-inflammatory response as summarized by the increase in macrophage polarization score upon treatment with CXCL16 (**Fig. 3B**). This macrophage score is an aggregate assessment of macrophage phenotype which is calculated by the ratio of pro-inflammatory (*Il1b*, *Il6*, *Mmp1b*) to anti-inflammatory (*Cd206*, *Il4*, *Il10*) gene expression (**Fig. S3A-B**). Interestingly, CXCL16 elicited a cell-type specific response as BMPCs treated with CXCL16 upregulated their expression of anti-inflammatory transcripts *Il10* and *Il1ra*, and the pro-inflammatory transcript *Il1b*, but *Il6* was unchanged (**Fig. 3C**).

**Fig. 3.**
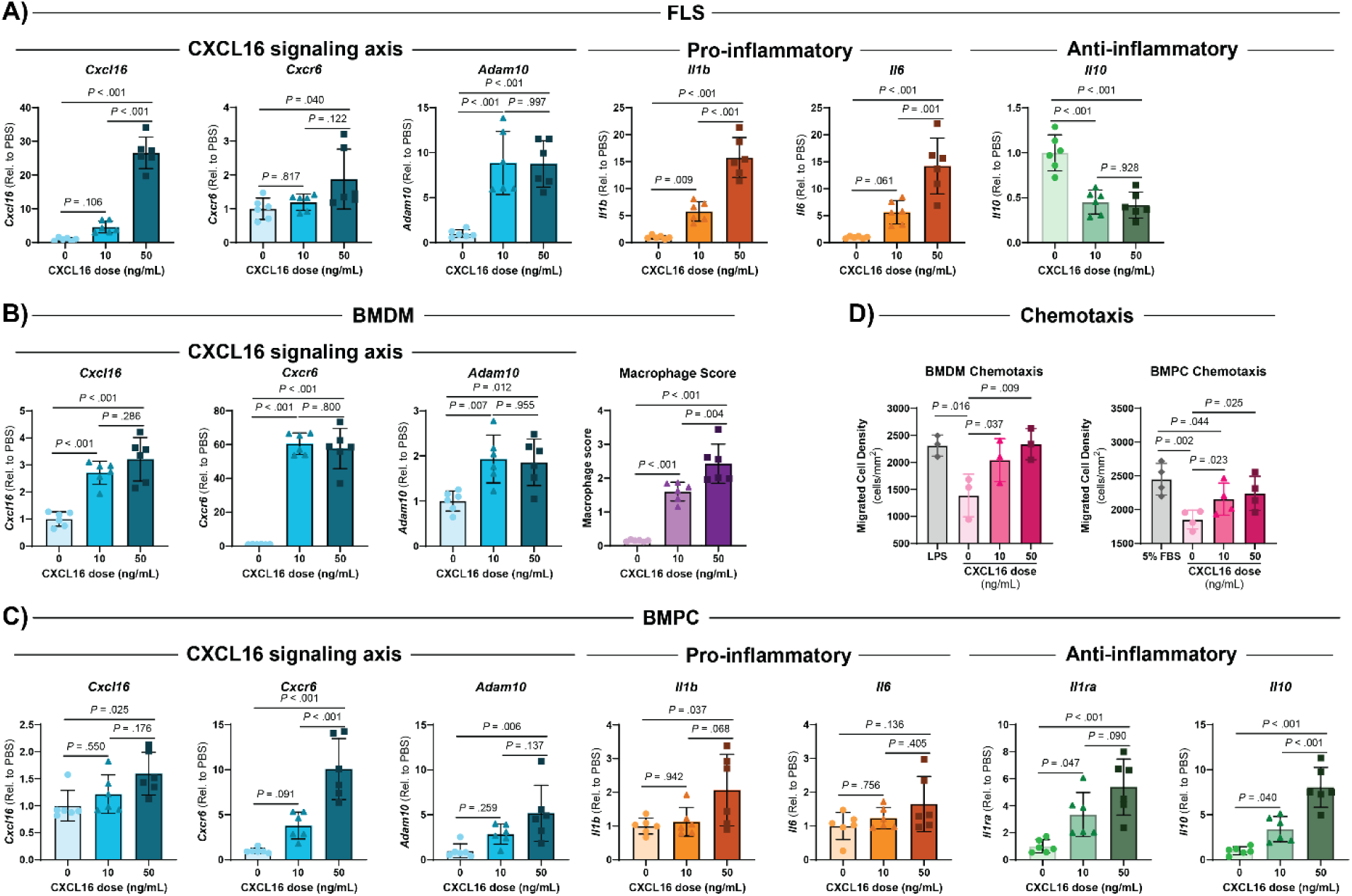
Downstream effects of CXCL16 *in vitro*. (**A**) In FLS, gene expression of the CXCL16 signaling axis, pro-inflammatory markers, and anti-inflammatory markers was measured by qPCR following stimulation with recombinant CXCL16 (n=6 per group). (**B**) In BMDMs, gene expression of the CXCL16 signaling axis and a macrophage score was measured by qPCR following stimulation with recombinant CXCL16 (n=6 per group). (**C**) In BMPCs, gene expression of the CXCL16 signaling axis, pro-inflammatory markers, and anti-inflammatory markers was measured by qPCR following stimulation with recombinant CXCL16 (n=6 per group). (**D**) Chemotaxis of BMDMs and BMPCs in response to CXCL16 was measured by migrated cell density (n=3 per group). All bars show mean ± SEM.

In addition to promoting cell-type specific inflammatory responses, CXCL16 contributed to its own feed forward signaling activation in FLS, BMDM, and BMPCs. CXCL16-treated FLS upregulated *Cxcl16*, *Cxcr6*, and *Adam10* expression (**Fig. 3A**). BMDMs exhibited similar responses, with increases in *Cxcl16*, *Cxcr6*, and *Adam10* expression after treatment with recombinant CXCL16 (**Fig. 3B**). BMPCs exhibited strong upregulation of *Cxcr6* with a less dramatic induction of *Cxcl16* and *Adam10* (**Fig. 3C**). In an *in vitro* migration assay, BMDMs and BMPCs also exhibited a dose-dependent chemotactic response to CXCL16 (**Fig. 3D**), corroborating prior studies on the chemotactic effects of CXCL16(*19–24*). Taken together, these *in vitro* findings demonstrate that CXCL16 induces pro-inflammatory effects in primary FLS and BMDMs but anti-inflammatory effects in BMPCs.

### Repeated intra-articular injections of CXCL16 induces knee hyperalgesia via CXCR6 and synovitis in healthy murine joints

On the basis of CXCL16 activating pro-inflammatory gene expression in synovial fibroblasts and BMDMs, we next wanted to assess whether CXCL16 is sufficient by itself to induce synovitis and joint disease *in vivo*. We injected either 5 µL of vehicle or 0.25 ng (low dose) or 25 ng (high dose) of recombinant murine CXCL16 into healthy, adult C57BL/6 knee joints for five consecutive days and performed knee hyperalgesia and histological assessments. Low and high dose CXCL16-injected limbs, but not vehicle-injected limbs, exhibited reduced knee withdrawal thresholds at 7d following the first injection, indicative of hyperalgesia (**Fig. 4A**). High dose CXCL16-injected limbs had a sustained reduction in knee withdrawal threshold up to 14d following the first injection, but this effect was not observed in low dose or vehicle-injected limbs (**Fig. 4A**). Histological evaluation demonstrated no change in the total post-traumatic osteoarthritis score, however there were significantly increased total synovitis scores in CXCL16-injected joints, driven by increased synovial lining hyperplasia and synovial fibrosis (**Fig. 4B-C**).

**Fig. 4.**
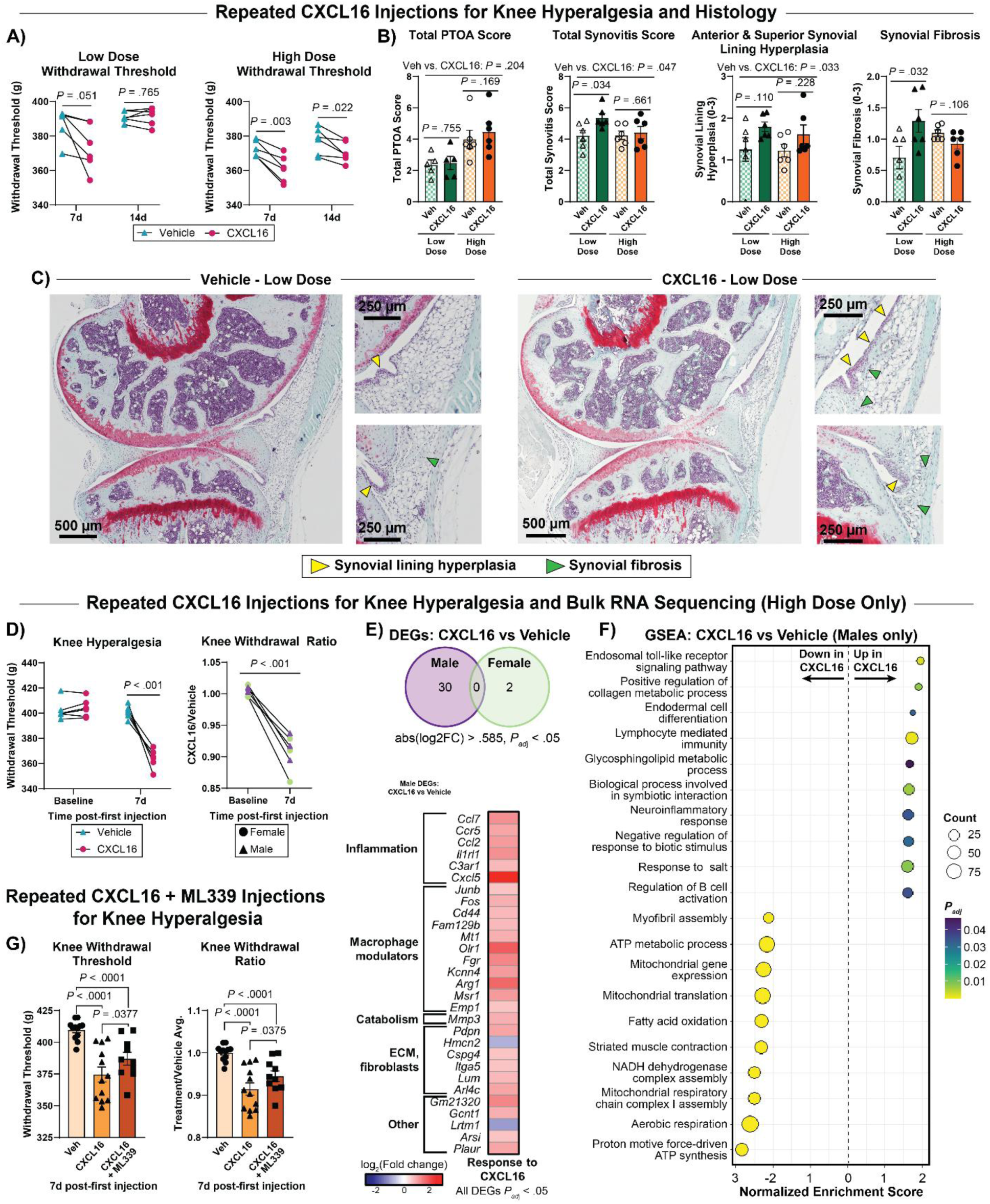
CXCL16-induced hyperalgesia and modulation of the synovial transcriptome. (**A**) Knee withdrawal threshold following repeated intra-articular injection of CXCL16 (either low dose or high dose) or vehicle in the contralateral side. Testing occurred at 7d and 14d post-first injection (n=6 per dose). (**B**) Histopathological scoring of limbs given repeated intra-articular injections of CXCL16 (either low or high dose) or vehicle in the contralateral side. Total PTOA score and total synovitis score are shown along with the synovitis subscores of anterior, inferior synovial lining hyperplasia and synovial fibrosis (n=5-6 mice per dose, n=3-12 images averaged together per limb). (**C**) Representative low dose CXCL16-injected limbs and vehicle-injected contralateral limbs. Yellow arrows – synovial lining hyperplasia. Green arrows – synovial fibrosis. (**D**) Knee withdrawal threshold and knee withdrawal threshold ratio (CXCL16/vehicle) following repeated intra-articular injection of CXCL16 (high dose only) or vehicle in the contralateral side (n=3 per sex). (**E**) Venn diagram of synovial bulk RNAseq showing differentially expressed genes (DEGs) between vehicle and CXCL16 injected joints in males compared to females (n=3 per group per sex, |log_2_FC| > .585, *P_adj_* < .05). Heatmap of male DEGs. Red indicates upregulated and blue indicates downregulated in CXCL16 injected joints relative to vehicle injected joints. (**F**) Gene set enrichment analysis of male synovia comparing vehicle vs CXCL16 injected joints. (**G**) Knee withdrawal threshold following repeated bilateral intra-articular injections of vehicle, CXCL16, or CXCL16 co-administered with ML339 (n=6 mice, i.e. n=12 limbs, per group). All bars show mean ± SEM.

To assess the molecular programs that underpin these pathological manifestations, we performed bulk RNA-seq of whole synovium from a separate cohort of mice treated with the identical vehicle or high-dose CXCL16 injection regimen as above. In this cohort, CXCL16-injected joints again exhibited reduced knee withdrawal thresholds 7d post-first injection (**Fig. 4D**). By RNA-seq at the same timepoint, we observed a highly sex-dependent perturbation of the synovial transcriptome due to CXCL16 administration (**Fig 4E**). Whereas female synovium exhibited near-absent CXCL16-induced differentially expressed genes (DEGs), male synovia upregulated cytokines, chemokines, and cytokine/chemokine receptors (e.g. *Ccl2, Ccl7, Cxcl5, Il1rl1, Ccr5*), genes of macrophage-relevant transcription factors and genes related to macrophage polarization (e.g. *Junb, Fos, Mt1, Arg1, Fgr*), and genes related to fibroblast activation and ECM turnover (*Pdpn, Mmp3, Itga5, Lum, Cspg4, Hmcn2*) (**Fig. 4E, Data file S2**). GSEA pathway analysis demonstrated that synovia from male CXCL16-treated joints had enriched pathways related to collagen production, immune/inflammatory response, and neuroinflammation (**Fig. 4F**). Taken together, these findings demonstrate that CXCL16 is sufficient by itself to induce arthritis-relevant disease manifestations in healthy joints *in vivo*, characterized by increased hyperalgesia, histological synovitis, and the induction of a synovial transcriptional program indicative of inflammation, lymphocyte and macrophage activation, ECM production, and fibroblast proliferation.

To interrogate the receptor mechanism by which CXCL16 induced pain, we co-administered a CXCR6-specific antagonist, ML339, in the same repeated injection study design. Male mice were administered five consecutive days of intra-articular vehicle, CXCL16 (200 ng), or CXCL16 (200 ng) + ML339 (1 mM) injections. CXCL16-injected joints again had reduced knee withdrawal thresholds and knee withdrawal threshold ratios as compared to vehicle injected limbs. This hyperalgesia-inducing effect was partially ablated by co-administration of ML339, as joints injected with combinatorial CXCL16 + ML339 had significantly higher knee withdrawal threshold and knee withdrawal ratio compared to joints injected with CXCL16 only (**Fig. 4G**).

### CXCL16 induces acute knee hyperalgesia via CXCR6

Joint pain is a complex physiological process orchestrated by multiple cell types, ligands, and receptors. A given mediator may agonize pain pathways via direct signaling, i.e. binding to joint nociceptor-bound receptors to directly alter nociception, or via indirect means, i.e. inducing the expression of other factors that directly bind nociceptors. Having demonstrated that CXCL16 is sufficient to induce knee hyperalgesia in a repeated-injection model, we sought to assess whether this at least partially occurs via direct interactions with joint-innervating nociceptors. We performed an acute hyperalgesia experiment in which recombinant CXCL16 (25 ng), or vehicle (sterile PBS) was injected into joints followed by knee withdrawal threshold measurements at acute timepoints post-injection (30 mins – 4 hrs) (**Fig. 5A**). Vehicle injection caused minor decreases in knee withdrawal threshold compared to baseline, however, CXCL16 significantly reduced withdrawal thresholds compared to vehicle by 30 mins post-injection in male mice, with a sustained effect until 7 days (**Fig. 5B-C, Fig. S4A-B**). Strikingly, despite female mice also exhibiting increased hyperalgesia after repeated injections in the prior experiment (**Fig. 4A, D**), female mice had absent hyperalgesia responses relative to vehicle in the acute experiment (**Fig. 5B-C**, **Fig. S4A-B**). The rapid responses observed in this experiment provide evidence of a direct nociceptive effect, i.e. direct interactions between CXCL16 and nociceptor-bound receptors(*37*, *38*).

**Fig. 5.**
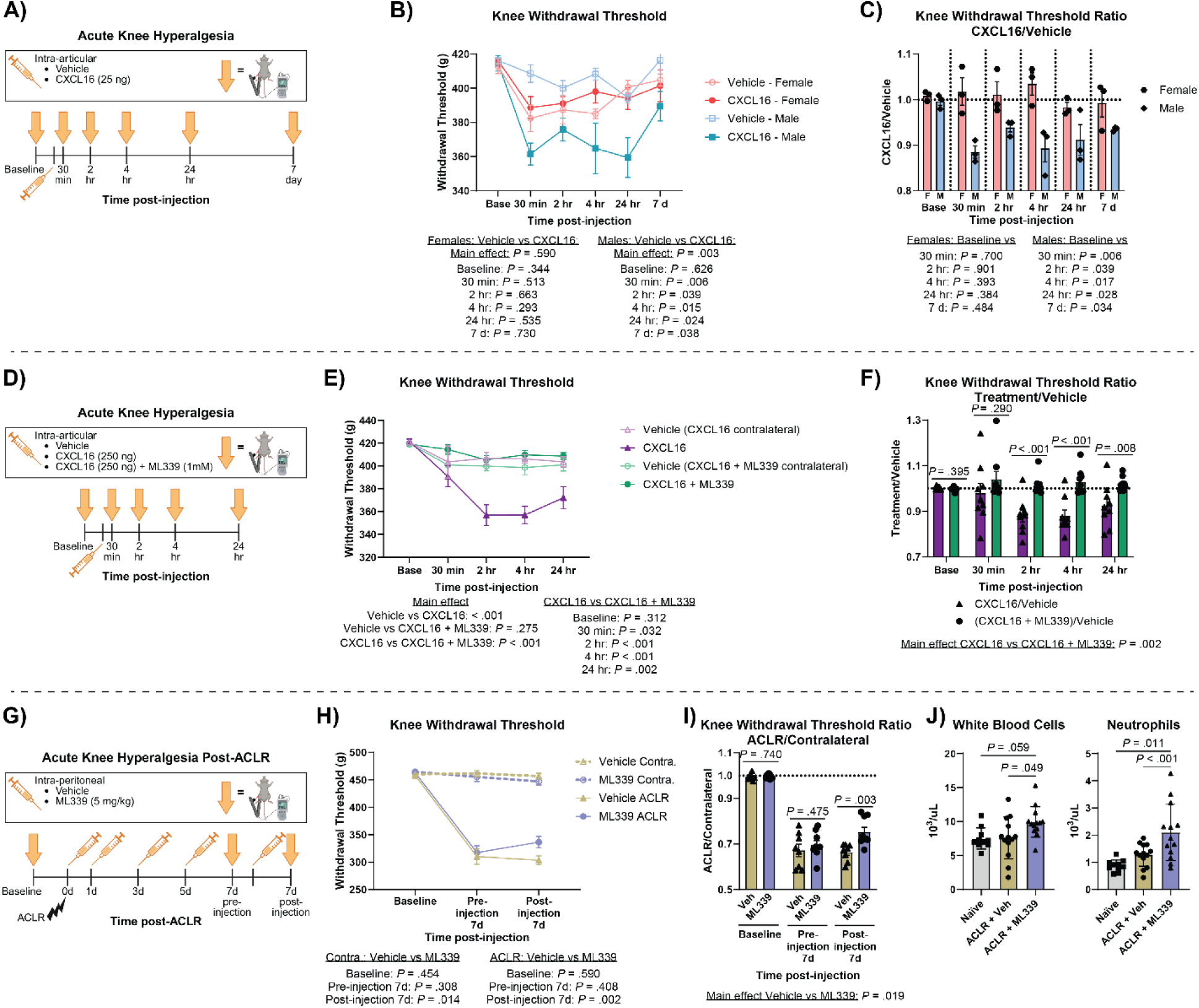
CXCL16-induced acute hyperalgesia via CXCR6. (**A**) Experimental design for intra-articular CXCL16 injection followed by acute knee hyperalgesia. Knee withdrawal threshold (**B**) and knee withdrawal threshold ratio (**C**) at acute timepoints following intra-articular injection of CXCL16 or vehicle in the contralateral joint (n=3 per sex). (**D**) Experimental design for intra-articular CXCL16 +/- ML339 injection followed by acute knee hyperalgesia. Knee withdrawal threshold (**E**) and knee withdrawal threshold ratio (**F**) at acute timepoints following intra-articular injection of CXCL16 or CXCL16 + ML339, and vehicle in the contralateral joint (n=9 per group). (**G**) Experimental design for intra-peritoneal ML339 injections and acute knee hyperalgesia post-ACLR. Knee withdrawal threshold (**H**) and knee withdrawal threshold ratio (**I**) pre- and post-intraperitoneal injection of vehicle or ML339 at 7d post-ACLR (n=8-9 per group). (**J**) Complete blood count analysis showing total number of circulating white blood cells and neutrophils from naïve mice and mice treated with vehicle or ML339 7d post-ACLR (n=10-13 per group). All bars show mean ± SEM.

We next wanted to ascertain whether the induction of acute hyperalgesia by CXCL16 is mediated by CXCR6. We conducted another acute hyperalgesia experiment in only male mice with a higher dose of CXCL16 (250 ng) and intra-articular co-administration of either vehicle or the CXCR6 antagonist ML339 (1mM) (**Fig. 5D**). Consistent with the prior experiment, CXCL16 induced acute hyperalgesia that persisted until 24 hrs post-injection, and co-administration of ML339 completely ablated this response (**Fig. 5E-F, Fig. S4C-D**). These findings demonstrate that CXCL16-induced hyperalgesia is mediated via CXCR6 binding.

### Systemic CXCR6 antagonism blocks joint injury-induced hyperalgesia

As joint injury induces the upregulation of multiple cytokines, chemokines, and neurotrophic mediators (**Fig. 1A**)(*32*) that orchestrate synovitis and hyperalgesia, we next wanted to assess whether CXCR6 antagonism mitigates injury-induced hyperalgesia. Male mice underwent ACLR and received 5x intra-peritoneal injections of ML339 (5 mg/kg) or vehicle followed by assessments of knee hyperalgesia and circulating immune cell populations (**Fig. 5G**). At the study endpoint, 7d post-ACLR, we also conducted an acute hyperalgesia experiment to measure any potential acute analgesic effects due to systemic injection. Whereas pre-injection withdrawal thresholds were comparable between treatment groups, ML339-treated mice had significantly higher withdrawal threshold ratio and absolute withdrawal threshold 2 hrs post-injection. (**Fig. 5H-I, Fig. S4E-F**). Complete blood counts further revealed that ML339-treated mice had higher circulating total white blood cell and neutrophil counts (**Fig. 5J**). These findings indicate that systemically administered ML339 acutely mitigates ACLR-induced knee hyperalgesia.

### CXCL16 induces rapid Ca^2+^ influx in dorsal root ganglia-derived nociceptive neurons via CXCR6

Given our observation that CXCR6 antagonism mitigated CXCL16-induced acute hyperalgesia as well as joint injury-induced hyperalgesia, we last sought to determine whether CXCL16 directly agonizes nociceptive neurons and whether this requires CXCR6. We isolated and cultured primary neurons derived from the dorsal root ganglia (DRG) of Na_V_1.8-Cre; GCaMP6s^fl/+^ mice, which express the fluorescent calcium indicator, GCaMP6s, in nociceptive neurons, marked by expression of the voltage-gated sodium channel 1.8 (Na_V_1.8)(*39*). In a time series experiment, cultured neurons were first treated with a negative control (vehicle only), followed by a single dose of 50 ng/ml CXCL16, with KCl used as a positive control (**Fig. 6A**). This was conducted in the presence or absence of 180 µM ML339 to antagonize CXCR6, and fluorescent signal was monitored and quantified as the change over baseline. CXCL16, but not vehicle, induced fluorescent responses, indicative of rapid neuronal Ca^2+^ signaling (**Fig. 6B-D**). Co-treatment with the CXCR6 antagonist ML339 completely ablated CXCL16-induced fluorescent responses (*P*<0.0001), with no effect on positive control responses to KCl (**Fig. 6B-D**). Taken together, these findings confirm that CXCL16 directly agonizes nociceptive neurons by inducing Ca^2+^ signaling, and this effect is abolished by pharmacological inhibition of the CXCR6 receptor.

**Fig. 6.**
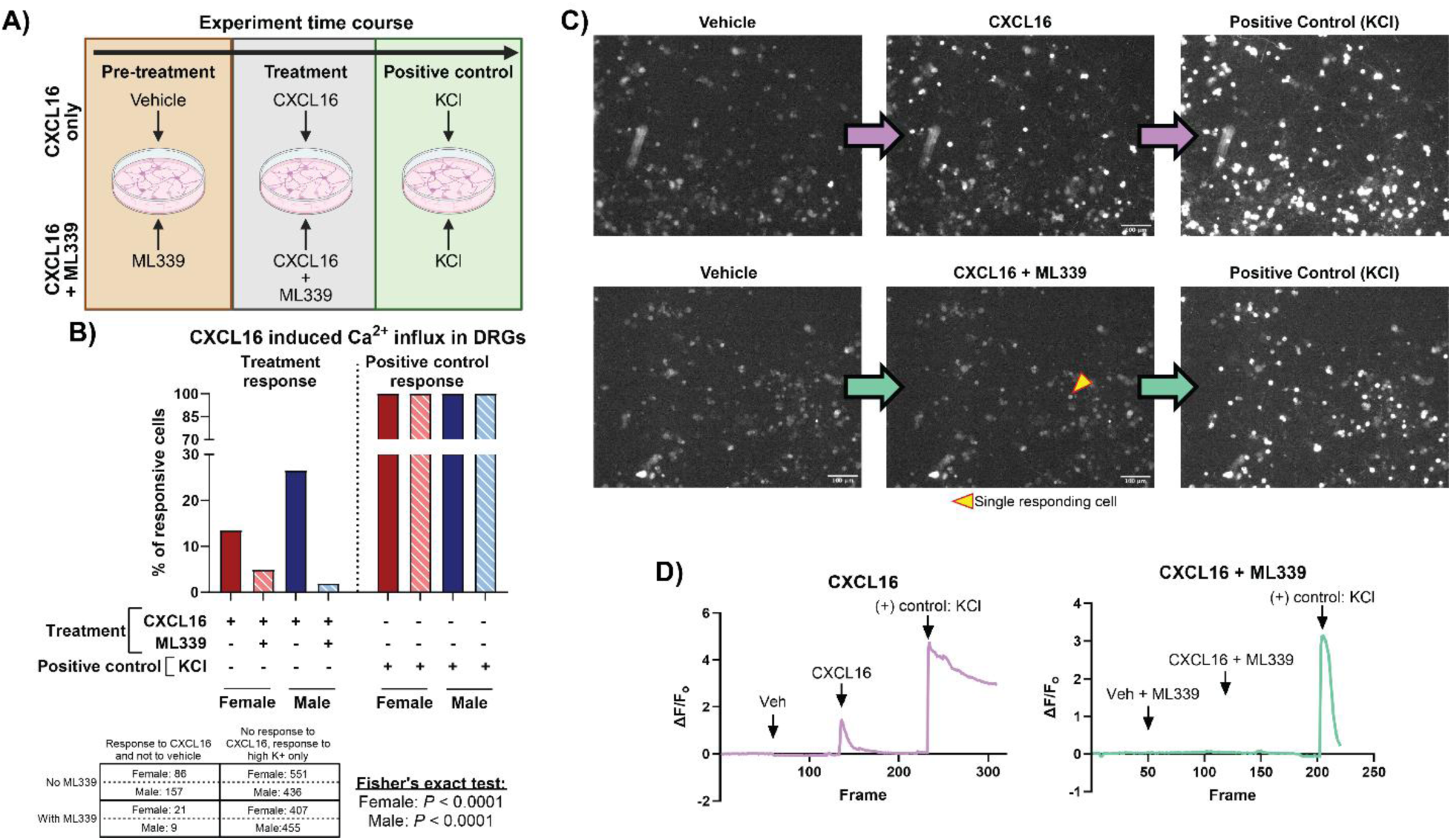
CXCL16-induced Ca^2+^ in Na_V_1.8 nociceptive neurons *in vitro*. (**A**) Experimental design: Murine DRG cultures were pre-treated with either vehicle or ML339, treated with CXCL16 or CXCL16 + ML339, and then followed by a positive control stimulus, KCl. (**B**) Quantification of percent responsive cells to CXCL16 stimulation in the presence and absence of CXCR6 antagonist, ML339. Total number of responsive cells is detailed for each sex. (**C**) Representative images of each stage of the experiment – pretreatment, treatment, positive control. (**D**) Representative line plots showing ΔF/Fo of a single responsive cell (responds to KCl) for each condition.

## DISCUSSION

This study elucidated the role of the CXCL16-CXCR6 axis in the context of post-traumatic osteoarthritis (PTOA), with a focus on its contributions to inflammation and nociception. Our findings demonstrate that joint injury rapidly activates gene and protein expression of CXCL16 and CXCR6, and that CXCL16 orchestrates synovial inflammation, pain-related behaviors and nociceptive signaling during PTOA progression. Using a translationally-relevant small-molecule antagonist against CXCR6, we confirmed that CXCR6 mediates CXCL16-induced joint hyperalgesia *in vivo* as well as CXCL16-induced Ca^2+^ signaling in DRG-derived nociceptive neurons *in vitro*.

To our knowledge, this is the first study to demonstrate the role of CXCL16 and its only known classical receptor, CXCR6, in peripheral nociception. Recently, Wang, et. al., demonstrated that CXCL16 expression and CXCL16 mRNA N^6^-methyladenosine modification was increased in rat sympathetic ganglia in a model of spared nerve injury (SNI)-induced neuropathic pain(*40*). They further showed that intra-DRG administration of CXCL16 induced mechanical allodynia in naïve rats while ablation of CXCL16 in sympathetic ganglia mitigated SNI-induced mechanical allodynia, establishing a role for CXCL16 in mediating neuropathic pain(*40*). Wang, et. al., also observed CXCR6 protein expression in rat DRG neurons(*40*), which suggests that CXCL16 can directly bind to DRG neurons via CXCR6. These findings support our premise that CXCL16 can induce knee hyperalgesia through direct binding to CXCR6+ nociceptive neurons. Of note, Wang, et. al did not empirically test whether CXCL16 operates via CXCR6, and our study did not measure CXCR6 expression by joint-resident nociceptors, which is an important future direction. However, using a small-molecule CXCR6 antagonist, we confirmed the receptor mechanism of CXCL16-induced effects in cultured DRG neurons *in vitro* and in CXCL16-induced joint hyperalgesia *in vivo,* providing the first mechanistic evidence for a CXCL16-CXCR6 binding interaction that promotes peripheral nociceptive signaling and pain-related behaviors.

CXCL16 has been shown to induce calcium mobilization in a CXCR6-expressing murine pre-B lymphoma cell line(*17*) and previous studies in rat and mouse DRGs have established the direct activation of Ca*^2+^* mobilization by other chemokines, including CCL2/MCP-1, CCL5/RANTES, CXCL1/fractalkine(*30*), and CXCL5(*31*). However, to date, CXCL16 has not been tested in this context. Our studies advance the understanding of the role of CXCL16-CXCR6 in calcium mobilization, as we demonstrated that CXCL16 can directly stimulate calcium signaling in mouse DRG neurons via CXCR6. Although a direct connection between CXCL16-CXCR6 and pain-related behaviors has not previously been established, CXCL16 has been connected to the nervous system through roles in neurotransmitter release and neuronal-glial cellular crosstalk(*41–43*). CXCL16 and CXCR6 have been shown to be expressed by astrocytes, microglia, and neurons in hippocampal cultures, indicating a potential for CXCL16-CXCR6 signaling(*42*). Di Castro, et. al, observed that CXCL16 promotes spontaneous GABA release in pyramidal neurons from the hippocampal CA1 area(*41*). Wang, et al. also showed that CXCL16 worked synergistically with norepinephrine to increase DRG hyperexcitability and exacerbate neuropathic pain(*40*). Our finding that CXCL16 induces acute knee hyperalgesia, as early as 30 minutes post-injection, suggests that CXCL16 is interacting directly with intra-articular nociceptors, but further studies are required to determine whether this hypothesized activation is due to modulation of Ca^2+^ influx, neurotransmitter release, or some other mechanism.

CXCL16 and CXCR6 have been established as inflammatory markers across multiple diseases including high-grade prostate cancer, which is considered an inflammation associated cancer(*44*). Lehrke, et. al., also suggest a pro-inflammatory role for CXCL16 in inflammatory bowel disease as CXCL16 was highly enriched in patients with Crohn’s disease or ulcerative colitis and the elevated CXCL16 levels correlated with level of C-reactive protein(*45*). Our studies also demonstrate a pro-inflammatory role for CXCL16, showing that CXCL16 elicits a pro-inflammatory response in synovium *in vivo* and in fibroblasts and macrophages *in vitro*. However, these responses were not tested alongside the CXCR6 antagonist ML339, and thus we cannot conclude whether pro-inflammatory effects were mediated by CXCR6. Further, we demonstrate mitigated CXCL16-induced pain following co-administration with ML339 in a repeated injection experimental design, but we did not evaluate whether co-administration with ML339 also mitigated the pro-inflammatory activation of the synovium that was observed in males. Future studies aim to address these limitations. In a cardiotoxin injection model of muscle injury, CXCL16 knockout mice had decreased macrophage infiltration but increased neutrophil infiltration to injured muscle as compared to controls(*46*). CXCL16 knockout mice also had decreased infiltration of Ly6C^hi^ monocytes and macrophages in the infarcted heart following myocardial infarction(*47*). These studies and our data support the function of CXCL16 as a macrophage chemoattractant across multiple tissue types, and our findings extend this knowledge by demonstrating its pro-inflammatory effects in synovial fibroblasts.

To the author’s knowledge, while the CXCR6 antagonist ML339 has been validated *in vitro*(*48*), *in vivo* use has been limited to a few cancer studies(*49*, *50*) and no study to date has used ML339 *in vivo* under any musculoskeletal health context. While systemic administration of the CXCR6 antagonist acutely ablated joint injury-induced hyperalgesia, it did not mitigate overall hyperalgesia development via disease modification at 7d post-ACLR. These findings indicate that the development of joint hyperalgesia following injury is not solely dependent upon CXCR6, but that CXCL16-CXCR6 at least partially mediate acute nociception. As CXCL16 is the only described ligand for CXCR6, we conclude that the injury-induced upregulation of CXCL16 in the joint, which we confirmed at the protein level in synovial fluid, is partially responsible for injury-induced hyperalgesia, which can be transiently blocked by targeting the CXCR6 receptor. The notion of synovium-derived CXCL16 ligand inducing knee hyperalgesia by binding to intra-articular nociceptors is supported by a meta-analysis of human osteoarthritic synovium. Newton, et. al, unbiasedly identified a putative synovium-DRG interactome focused on neuro-immune/neuro-inflammatory pathways, of which synovium-derived CXCL16 was predicted to interact with nociceptor-bound CXCR6(*51*).

Our findings should be interpreted in light of several limitations. The sex-specific differences in the synovial transcriptome and acute hyperalgesia observed in response to intra-articular CXCL16 injection highlight an area needing further investigation. Such studies should address whether this is related to a dose dependency. This will allow future refinement of treatment approaches that account for sex as a biological variable. Furthermore, the efficiency of *in vivo* CXCR6 antagonism by ML339 is unclear, and residual CXCR6-mediated signaling by incomplete receptor inhibition may compensate certain responses, which should be addressed by studies employing genetic knockouts of CXCL16 and CXCR6. The magnitude of contribution of direct CXCL16 binding to intra-articular nociceptors versus secondary stimulation of nociceptors via CXCL16-driven immune cell activation to *in vivo* CXCL16-dependent pain regulation requires further investigation. While our study provides a comprehensive examination of the role of CXCL16 within murine models, there is a need to validate these findings in larger animal models, and human synovial cells and DRGs. Moreover, our study focused mainly on the acute phases of PTOA, with limited assessments of the long-term effects of CXCL16 activation or CXCR6 inhibition. Chronic pain and progressive joint degeneration are characteristic of PTOA, and thus longer-term studies are needed to understand whether the analgesic effects of CXCR6 inhibition are sustained throughout PTOA progression.

In summary, our study unveils the CXCL16-CXCR6 axis as a novel mediator of joint inflammation and nociception in PTOA while also establishing pharmacological CXCR6 antagonism as a potential therapeutic strategy to treat joint trauma-related inflammation and pain.

## MATERIALS AND METHODS

### Study design

The objectives of this study were to: 1) Evaluate the CXCL16 signaling axis in the context of PTOA; 2) establish key mediators and downstream effectors of CXCL16; 3) identify the sufficiency of CXCL16 in driving synovial inflammation and joint nociception, and finally, 4) establish a receptor mechanism through which CXCL16 exerts its effects. To explore how CXCL16 expression is altered following murine joint injury and in human OA synovium, we mined bulk and single cell RNAseq datasets and RNA microarrays from mouse and human synovium. Flow cytometry was used to identify CXCL16+ and CXCR6+ cells in healthy and injured mouse synovium, and ELISA was used to measure CXCL16 protein in synovial fluid. *In vitro* studies examined the effects of pro-inflammatory cytokines and CXCL16 treatment in murine joint-relevant cell types. The sufficiency of CXCL16 to induce synovial inflammation and joint nociception *in vivo* was evaluated by repeated joint injections with recombinant CXCL16 in naïve mice, followed by knee hyperalgesia testing, histological evaluation, and bulk RNA sequencing of synovium. The receptor mechanism by which repeated intra-articular dosing of CXCL16 induces knee hyperalgesia was assessed by co-administration of CXCL16 and a CXCR6 antagonist, ML339. To demonstrate whether CXCL16 was directly agonizing nociceptors to induce pain, mice were administered a single intra-articular injection of CXCL16 followed by hyperalgesia measurements as early as 30 mins post-injection. This was repeated with co-administration of ML339 to establish CXCR6 as the receptor mediating this effect. The role of CXCL16 in driving joint injury-induced pain was tested by administering multiple intra-peritoneal doses of ML339 following noninvasive joint injury in mice and performing knee hyperalgesia assessments longitudinally. Finally, to assess whether CXCL16 can directly activate calcium signaling in nociceptive neurons, we performed *in vitro* calcium imaging of primary DRG-derived neurons from Na_V_1.8-Cre; GCaMP6s^fl/+^ mice following treatment with CXCL16, with or without CXCR6 antagonist co-treatment. Mice were randomly assigned to groups within an experiment. A blinded operator performed all knee hyperalgesia testing and histopathologic scoring. Sample sizes are detailed in the figure caption and statistical analyses are described in the *Data analysis and statistics* methods section.

### Animals and noninvasive ACL rupture

All animal studies were approved by the institutional animal care and use committee. Male and female C57BL/6 mice aged 12-15 weeks (Jackson Laboratories) were used throughout. To induce joint injury and PTOA, we employed a noninvasive mechanical loading-based anterior cruciate ligament rupture (ACLR) model(*32*, *36*) (**Suppl. Materials and Methods**). Limbs from either Sham mice (no loading, with anesthesia and analgesia only) or unloaded contralateral limbs from injured mice served as controls. Naïve male and female Na_V_1.8-Cre; GCaMP6s^fl/+^ mice(*39*) older than 12 weeks were used to obtain primary DRG cultures for *in vitro* Ca^2+^ imaging experiments.

### Bulk RNA sequencing, single-cell RNA-sequencing, RNA microarray published data mining

Existing, published bulk RNAseq data generated by our group(*32*) was re-analyzed to assess injury-induced gene expression of all CCL and CXCL family chemokines and their receptors in Sham, ACLR 7d, and ACLR 28d male and female synovia (**Suppl. Materials and Methods**).

To assess expression patterns of Cxcl16, Cxcr6, and Adam10 in synovium, existing published scRNAseq data from mouse and human OA synovium was analyzed using R. Mouse scRNAseq previously published in our lab(*35*) was mined to examine synovial cell types expressing *Cxcl16*, *Cxcr6*, and *Adam10*. Feature plots showing the expression of the genes of interest across the all cells object, which contains integrated Sham, 7d ACLR, and 28d ACLR synovial cells were generated using the *Shiny* R package.

A meta-analysis of human OA synovium bulk RNAseq and RNA microarray datasets was performed using weighted Limma-Voom. Data from these is given as log_2_(fold change) and -log10(*P_adj_*). Unadjusted *P*-values from individual studies were combined across datasets into a composite *P*-value using the weighted *Z*-test, with a BH (e.g. FDR) correction for multiple comparisons applied following combination. This test is directional, so a positive change in one study and negative change in another cancel out rather than combining. Composite data is presented only as a signed -log_10_(*P_adj_*), since log_2_(fold change) cannot be validly combined between RNAseq and RNA microarray datasets. For genes which were missing from some microarrays, only datasets which contained these genes were included in the composite *P*-value calculation.

### Synovial fluid collection and protein quantification

Synovial fluid was collected from joints as previously described(*36*, *52*). In brief, following euthanasia, a small, consistently-sized alginate sponge was inserted into the joint space through a small incision made in the suprapatellar capsule. Synovial fluid-soaked sponges were flash frozen and stored at -80C. Following thawing, sponges were digested via alginate lyase, and digestates were assayed for CXCL16 content by ELISA (Invitrogen: EMCXCL16). Total protein was quantified in the same samples via Pierce™ Bradford Plus Protein Assay Kit (ThermoFisher Scientific, Cat. #23236). CXCL16 concentrations were normalized to total protein.

### Flow cytometry

Whole synovium, inclusive of Hoffa’s fat pad, was dissected and enzymatically digested to yield single-cell suspensions using Liberase TM, Collagenase IV, and DNaseI(*35*, *36*). Each sample consisted of one male and one female synovium pooled at the time of dissection (n=3 ACLR, n=3 contralateral). After digestion, single cell suspensions were stained (**Suppl. Materials and Methods**). Flow cytometry was performed on a BD LSRFortessa and analysis was performed in FlowJo v10 software (BD/TreeStar).

### Tissue gene expression via RT-qPCR

Dissected joint tissue were homogenized in TRIzol, RNA was isolated, and qPCR was performed to evaluate the expression of target genes (**Suppl. Materials and Methods**). Data was normalized to housekeeping genes and reported further normalized to experimental controls using the 2^-ΔΔCT^ method.

### Primary cell isolation and in vitro treatments

Primary fibroblast-like synoviocytes (FLS) from knee synovia, bone marrow-derived macrophages (BMDMs), and bone marrow-derived mesenchymal progenitor cells (BMPCs) were isolated from naïve adult male and female C57BL/6 mice, as previously described(*35*) (**Suppl. Materials and Methods).**

To ascertain upstream activators of *Cxcl16* and related gene expression, FLS, BMDMs, and BMPCs were treated with PBS (control), TNF-α (10 ng/mL), or IL-1β (10 ng/mL) for 48 hours (FLS, BMPCs) or 8 hours (BMDMs) and then harvested and processed for RT-qPCR. Conditioned media from FLS was also collected for CXCL16 protein quantification via ELISA. To assess the potential functional role of CXCL16, FLS, BMDMs, and BMPCs were treated with PBS (control) or CXCL16 (10ng/mL or 50 ng/mL) for 48 hours (FLS, BMPCs) or 8 hours (BMDMs) and then harvested and processed for RT-qPCR. All *in vitro* experiments were conducted in n=4-8 biological replicates, i.e. cell isolations from unique mice (**Suppl. Materials and Methods)**.

### ATDC5 cell culture

Following differentiation (**Suppl. Materials and Methods**), chondrogenic ATDC5s were treated with PBS (control), TNF-α (10 ng/mL), or IL-1β (10 ng/mL) for 48 hours, followed by gene expression studies. Briefly, ATDC5s were lysed with Trizol and frozen at -80**°**C until RNA isolation and gene expression studies by RT-qPCR. All experiments were performed in n=4 replicates.

### In vitro chemotaxis of BMDMs and BMPCs

BMDMs and BMPCs were separately seeded into transwell insert membranes and combined with a transwell chamber containing either media with vehicle, a positive control (LPS for BMDMs, 5% FBS for BMPCs), or recombinant mouse CXCL16 (10 ng/mL or 50 ng/mL). Cells were allowed to migrate for three days prior to quantification of migrated cell density (**Suppl. Materials and Methods**).

### Intraarticular and systemic treatments

Joint injections were administered to the knees of naïve C57Bl/6 mice. All injections were performed using 33G needles and microsyringes (Hamilton Company). To assess the disease-promoting effects of CXCL16 in healthy joints, CXCL16 (R&D Systems, Cat. #503-CX) (0.25 ng or 25 ng in 4 µL of sterile PBS) was injected for five consecutive days into one knee joint, and vehicle was injected in the contralateral joint (n=3 male, n=3 female per dose). Knee hyperalgesia testing occurred at 7d and 14d post-first injection. At 14d post-first injection hindlimbs were harvested for histological evaluation of PTOA and synovitis severity. A separate group of mice (n=3 male, n= 3 female) underwent a similar regiment of 5 consecutive intra-articular injections of CXCL16 into one knee joint (25 ng in 4 µL of sterile PBS with .1% BSA) and vehicle (sterile PBS with .1% BSA) into the contralateral. A baseline measurement of knee hyperalgesia was taken pre-injection then knee hyperalgesia and synovial harvests for bulk RNA sequencing occurred 7d post-first injection. In another cohort, mice received 5 consecutive days of bilateral intra-articular injections of either vehicle (sterile PBS with 0.69% DMSO), CXCL16 (200 ng in 5 µL sterile PBS with 0.69% DMSO), or CXCL16 co-administered with CXCR6 antagonist, ML339 (MedChemExpress, Cat. #HY-122197) (200 ng CXCL16 in 5 µL sterile PBS with 1mM ML339 and 0.69% DMSO). Knee hyperalgesia was performed at baseline and 7d post-first injection (n=6 males/group).

For the first acute hyperalgesia experiment in naïve mice, mice received either a single CXCL16 (25 ng in 4 µL of sterile PBS with 0.1% BSA) or vehicle (sterile PBS with 0.1% BSA) injection into a single joint (n=3 male, n=3 female). Knee hyperalgesia occurred at baseline and 30 min, 2 hr, 4 hr, 24 hr, and 7d post-injection (**Fig. 5A**). Mice were divided into two groups for the second acute hyperalgesia experiment in naïve mice (n=9 males/group). One group received an intra-articular injection of CXCL16 (250 ng in 6 µL sterile PBS containing 0.08% BSA and 0.69% DMSO) in one knee and vehicle (sterile PBS containing 0.08% BSA, and 0.69% DMSO) in the contralateral. The other group received an intra-articular injection of CXCL16 co-administered with CXCR6 antagonist, ML339, (250 ng CXCL16 in 6 µL sterile PBS containing 1mM ML339, 0.08% BSA, and 0.69% DMSO) in one knee and vehicle (sterile PBS containing 0.08% BSA, and 0.69% DMSO) in the contralateral. Knee hyperalgesia occurred at baseline and 30 min, 2 hr, 4 hr, and 24 hr (**Fig. 5D**). To assess whether ML339 mitigates ACLR-induced effects, injured mice (n=9 males/group) received repeated intraperitoneal injections of either vehicle (corn oil containing 1% DMSO) or ML339 (5 mg/kg of 1.45 mM ML339 in corn oil containing 1% DMSO). Knee hyperalgesia occurred at baseline, 7d post-ACLR prior to injection, and 7d post-ACLR at 2 hrs post-injection (**Fig. 5G**). All mice were randomly assigned to treatment groups within each experiment.

### Knee hyperalgesia testing

Knee hyperalgesia testing was performed by a blinded operator (LL) using a Randall-Selitto device as described previously(*32*, *36*). Briefly, pressure was applied to the medial aspect of the knee and the knee withdrawal threshold was recorded when the mouse gave a response – a struggle or vocalization. Three measurements were recorded for each limb and averaged together to obtain the knee withdrawal threshold per limb.

### Histological analysis

Whole hindlimbs were processed, stained with Safranin O/Fast Green (SafO), imaged (**Suppl. Materials and Methods**), and qualitatively graded for OA and synovitis severity using updated and extended versions (**Table S3-S4**) of an established grading scheme(*53*, *54*). All grading was performed in a blinded fashion (LL). Scores were averaged across all slides to calculate a limb average, and limbs were averaged to calculate group scores.

### Bulk RNA-sequencing and bioinformatic analysis

RNA was isolated from Trizol homogenates of whole synovium, inclusive of Hoffa’s fat pad, then sequenced and quality control was performed (**Data file S1**). All Bulk RNA-sequencing data analysis including quality control, gene expression analysis, and pathway analysis was performed in RStudio (R v4.4.0, RStudio v2024.04.1). Differential gene expression analysis was performed using DESeq2(*55*). Pathway analysis was performed using the *clusterProfiler*(*56*) implementation of fast gene set enrichment analysis (GSEA), using the GO: Biological Processes gene ontology terms list (**Suppl. Materials and Methods**).

### Complete blood counts

Complete blood count (CBC) analysis was performed using 50-100 µL of blood, obtained via cardiac puncture at the time of harvest.

### Primary DRG neuron culture and Ca^2+^ imaging

Primary DRG cultures were generated from Na_V_1.8-Cre; GCaMP6s^fl/+^ mice as previously described(*57*). These mice express the fluorescent calcium indicator, GCaMP6s, in nociceptive neurons, marked by expression of the voltage-gated sodium channel 1.8 (Na_V_1.8). L3-L5 DRGs were collected and pooled from 12 wk-old mice, and male and female cultures were generated separately. Four DRGs were used for every coverslip plated. Cultures were maintained at 37℃, treated with CXCL16 with or without pre-treatment with a CXCR6 antagonist, ML339 (180 µM), and fluorescent imaging was performed with Zeiss Zen software to measure Ca^2+^ signaling (n=2-5 well replicates per sex/treatment) (**Suppl. Materials and Methods**). KCl was used as a positive control.

Fluorescent imaging of the calcium sensor was performed at each step of the stimulations and a Fiji macro was used to create videos for calculating (Ft-Fo) / Fo, termed “ΔF/Fo”. Regions of interest (ROIs) were circled around individual neurons in Fiji based on responses to KCl and ΔF/Fo was recorded for each ROI. Graphs for each ROI were created in R or in Python. Each graph and video file were manually checked to determine responses for each neuron. Neurons that responded to vehicle were excluded from further analysis.

### Statistical analysis

SPSS (v27, IBM, Armonk, NY) and Prism 10.0 (Graphpad, San Diego, CA) were used for statistical analyses. Paired t-tests compared contralateral and ACLR groups for synovial fluid ELISA and synovial tissue qPCR and flow cytometry analyses. Ordinary one-way ANOVA with Tukey’s multiple comparison evaluated treatment differences for *in vitro* cytokine and CXCL16 treatments. Repeated measures one-way ANOVA was used to analyze *in vitro* chemotaxis experiments. Paired t-tests compared knee hyperalgesia threshold and knee withdrawal threshold ratio between vehicle and CXCL16 injected limbs. Paired t-tests within each dose and across both doses of CXCL16 compared histological parameters between vehicle injected and CXCL16 injected limbs. Two-way ANOVA was used to compare knee withdrawal threshold and knee withdrawal threshold ratio between vehicle, CXCL16, and CXCL16 + ML339 injected limbs at baseline and 7d post-injection (factors: timepoint, group). Linear mixed effects models were used to compare knee withdrawal threshold in acute hyperalgesia experiments (**Suppl. Materials and Methods**). Fisher’s exact tests within each sex were used to determine associations between the proportion of cells responsive to CXCL16 or to positive control only, in the presence or absence of ML339. *P* values below 0.05 were considered significant throughout.

### List of Supplementary Materials

Materials and Methods

Fig. S1. Synovial gene expression of chemokine ligands and chemokine receptors is perturbed in mouse and human OA.

Fig. S2. qPCR corroboration of CXCL16 signaling axis expression and flow cytometry supplement

Fig. S3. CXCL16 treatment *in vitro* – individual expression of macrophage score genes.

Fig. S4. Acute knee hyperalgesia supplement.

Fig. S5. Bulk RNAseq PCA plot.

Table S1. Antibodies.

Table S2. Gene primers for qPCR.

Table S3. PTOA severity scoring.

Table S4. Synovitis severity scoring.

Data file S1. Synovial bulk RNA-seq – quality control.

Data file S2. Synovial bulk RNA-seq – DEG lists.

## Acknowledgements

The authors would like to acknowledge the Michigan Integrative Musculoskeletal Health Core Center (MiMHC) which is supported by P30AR069620 from NIH/NIAMS for processing samples for histology and for use of their microscopy equipment. We also wish to acknowledge the University of Michigan Biomedical Research Core Facilities, specifically the flow cytometry core, which is a part of the Medical School Office of Research, for use of their flow cytometer. We would like to thank the Chicago Center on Musculoskeletal Pain (C-COMP) funded by P30AR079206 for technical assistance with calcium imaging.

## Funding

National Science Foundation Graduate Research Fellowship grant DGE 2241144 (LL)

National Institutes of Health K99 grant K99AR081894 (AJK)

Pioneer Fellowship from the University of Michigan (AJK)

Orthopaedic Research Society Scientific Network Award (TM, REM)

F31AR083277 (NSA)

R01AR077019 (REM)

R01AR077019-03S1 (REM)

R01AR080035 (TM)

Department of Orthopaedic Surgery, University of Michigan (TM)

## Author contributions

Conceptualization: LL, SR, REM, TM

Methodology: LL, SR, AJK, MDN, PMR, REM, TM

Investigation: LL, SR, AJK, AM, MDN, RFB, IS, SCH, SGN, PMR, NSA

Visualization: LL, AM, MDN, IS

Funding acquisition: LL, NSA, REM, TM

Writing – original draft: LL, TM

Writing – review & editing: LL, AJK, MDN, NSA, REM, TM

## Data and materials availability

Bulk RNA-seq data will be publicly available on GEO and all other data are available in the main text or the supplementary materials.

