## Supplemental materials and methods for "CXCL16 mediates nociception and inflammation in murine post-traumatic osteoarthritis"

*Animals and noninvasive ACL rupture*

Under isoflurane anesthesia (5% induction, 2.0-2.5% maintenance), mice were positioned prone on a custom fixture, with the right knee in 100° of flexion. Following preconditioning, a rapid 1.5 mm compressive displacement was applied to the tibia using a mechanical testing system (ElectroForce 3300AT, TA Instruments, New Castle, DE, USA or CellScale Univert S2, CellScale, Waterloo, Ontario, Canada) to induce a closed, isolated ACL rupture. For analgesia, mice were given a single injection of subcutaneous carprofen (5 mg/kg). Following injury, mice were allowed to continue *ad libitum* cage activity in a 12-hr light/dark facility. Mice were euthanized via $CO_2$ asphyxia.

*Bulk RNA sequencing, single-cell RNA-sequencing, RNA microarray published data mining*

Bulk RNAseq: Muscle contamination and PCA of read counts was used to identify outliers, resulting in 4 out of 60 samples excluded from subsequent analyses. Following removal of lowly-expressed genes (FPKM < 1), *DESeq2* was used for differential-expression analysis. Unpaired analyses were performed to compare between conditions (Sham vs 7d ACLR, Sham vs 28d ACLR). The *EnhancedVolcano* R package was used to create independent volcano plots of chemokine ligands and receptors. Cutoffs of $P < 0.05$ and $|\log_2 FC| > 0.585$ are shown with dotted lines.

*Flow cytometry*

After digestion, cells were incubated in PBS with eFluor660 Fixable Viability Dye (eBioscience, 1:1,000) for 30 min at 4$^{o}$C in the dark. Non-specific antibody binding was minimized by staining cells with mouse FcX TruStain PLUS (Biolegend, 1:1,000). Surface marker staining was then performed using a cocktail of fluorescently conjugated antibodies in FACS buffer (PBS, 2% FBS, 1mM EDTA), to identify major synovial cell types. Next, intracellular staining for CXCL16 was undertaken after fixation and permeabilization of cells using the Biolegend Cyto-Fast Fix/Perm Buffer Set. Antibody details are found in **Table S1**. Unstained cells, single-stained controls, and fluorescent-minus-one (FMO) controls were included for compensation setup and establishing positive and negative gating.

*Tissue gene expression via RT-qPCR*

Dissected joint tissues were snap-frozen in Trizol and stored at -80C until analysis. Upon thawing, samples were homogenized in Trizol using an orbital homogenizer (Precellys Evolution, Bertin Instruments, Rockville, MD, USA) and bead-containing homogenization tubes (Hard tissue homogenizing CK28, Precellys). RNA was isolated via chloroform phase separation and converted to cDNA (High Capacity cDNA Reverse Transcription Kit, Applied Biosystems, CA, USA). cDNA was combined with SYBR Green master mix (Applied Biosystems, MA, USA) and relevant gene primers (**Table S2**) for RT-qPCR with a Bio-Rad T100 Thermal Cycler (Bio-Rad, Berkeley, CA, USA).

*Primary cell isolation and in vitro treatments*

Primary FLS were isolated from naïve adult C57BL/6 knee synovium by enzymatic digestion in Liberase, Collagenase IV, and DNaseI for 30 minutes at 37°C. FLS were expanded in DMEM containing 10% FBS and 1% penicillin-streptomycin. Passage 3-4 cells were used in all experiments. Bone marrow-derived macrophages (BMDM) were isolated from naïve adult C57BL/6 mice. Bone marrow was centrifuged out of long bones and cultured in DMEM, 10% FBS, 20ng/mL m-CSF for 7 days. Bone marrow-derived mesenchymal progenitor cells (BMPC) were isolated by culturing mouse bone marrow in complete MSC culture medium (αMEM, 20% FBS) for 7 days, allowing BMPCs to proliferate, and then BMPCs were seeded into experimental wells at P1.

To ascertain upstream activators of *Cxcl16* and related gene expression, FLS, BMDMs, and BMPCs were seeded in 12 well plates (100,00- cells/well), serum starved overnight, and exposed to serum-free media containing PBS (control), TNF-α (10 ng/mL), or IL-1β (10 ng/mL) for 48 hours (FLS, BMPCs) or 8 hours (BMDMs). Following treatment, cells were lysed with Trizol and stored at -80°C, followed by RNA isolation and gene expression studies by RT-qPCR. Conditioned media from FLS was also collected and frozen at -80°C for downstream protein quantification of CXCL16 via ELISA. To assess the potential functional role of CXCL16, FLS, BMDMs, and BMPCs were similarly seeded in 12 well plates (100,00 cells/well), serum starved overnight, and exposed to serum-free media containing PBS (control) or CXCL16 (10ng/mL or 50 ng/mL) for 48 hours (FLS, BMPCs) or 8 hours (BMDMs). Again, following treatment, cells were lysed with Trizol and stored at -80°C, followed by RNA isolation and gene expression studies by RT-qPCR.

*ATDC5 cell culture*

ATDC5s were cultured and expanded in (Invitrogen DMEM Cat. #11965092, with 1% anti-fungal/anti-bacterial, 5% FBS). Prior to any treatment, ATDC5s underwent 21-day chondrogenic differentiation in (Invitrogen DMEM Cat. #11995065, with 40 μg/mL L-proline, 1X ITS premix, 1X L-glutamine, 1% anti-fungal/anti-bacterial, 40 μg/mL AA2P, 100 nM dexamethasone, 10 ng/mL TGF- β1) with media changes occurring every 3 days.

*In vitro chemotaxis of BMDMs and BMPCs*

BMDMs and BMPCs were serum starved overnight and seeded (150,000-200,000 cells/well) into transwell insert membranes (8μm pore size) in 0.5% FBS-containing media. The bottom transwell chamber contained either media containing vehicle, a positive control (LPS for BMDMs, 5% FBS for BMPCs), or recombinant mouse CXCL16 (10 ng/mL or 50 ng/mL). The transwell chambers were combined and cells were allowed to migrate for 3 days at 37°C. Following migration, the top of the membrane was carefully wiped, and cells on the bottom membrane were fixed in 70% ethanol, stained with Hoechst solution (1:2500 dilution), rinsed with PBS, and mounted on a slide for imaging and analysis. Ten evenly sized fields of view were obtained via fluorescent microscopy, and the number of cells per view was quantified using MATLAB. Migration experiments were conducted across n=3-4 biological replicates, i.e. unique cell isolations.

*Histological analysis*

Whole hindlimbs were skinned, fixed in 10% neutral-buffered formalin for 48 hrs, then decalcified in 10% EDTA for 14 days, and stored in 70% ethanol prior to being processed and embedded in paraffin. 5-μm sagittal sections of the medial knee compartment were collected. Slides were stained with Safranin O/Fast Green (SafO), imaged (Eclipse Ni E800 with DS-Ri2 camera, Nikon, Tokyo, Japan), and qualitatively graded for OA and synovitis severity (**Table S3-S4**).

*Bulk RNA-sequencing and bioinformatic analysis*

RNA was isolated and purified from Trizol homogenates of whole synovium, inclusive of Hoffa's fat pad, using a RNeasy® Mini Kit (QIAGEN, Hilden, Germany). Whole mRNA was sequenced using polyA capture (150 bp paired-end reads, >40M reads/sample) on a NovaSeq 6000 sequencer (Illumina Inc., San Diego, California, United States). Quality control of sequencing data confirmed high quality RNA (RIN > 7) and successful, accurate sequencing in all samples (**Data file S1**).

Raw read counts were filtered to exclude non-protein-coding genes, sex-linked genes. Lowly-expressed genes were also removed, with only genes with FPKM > 1 and > 10 total reads in at least $n$=3 samples retained. Outliers were assessed via principal component analysis (PCA), and significant statistical outliers were not detected (**Fig. S5A**). Differential gene expression analysis was performed using DESeq2 (*54).* A paired study design was employed, with sex as the between-subject factor and treatment as within-subject factor. The effect of CXCL16 treatment was assessed *within* each sex, and independent ("main") effects of sex (*independent* of treatment) and CXCL16 treatment (*independent* of sex) were also assessed. DEGs were defined as $P_{\mathrm{adj}} < 0.05$ with a fold-change filter of $|\log_2 FC|>0.585$ (i.e. >|1.5|-fold change). Gene expression heatmaps were generated by calculating the z-score from the raw count data for DEGs by subtracting the row mean from each value and dividing by the row standard deviation. The *pheatmap* R package was used for visualization of the z-score heatmaps.

Pathway analysis was performed using the *clusterProfiler (55)* implementation of fast gene set enrichment analysis (GSEA), using the GO: Biological Processes gene ontology terms list. For each comparison, all genes were assigned signed gene rankings according to the formula sign($\log_2$FC) * $P$-value (based on differential-expression analysis), sorted in descending order, and submitted to GSEA analysis via the *gseGO* function. GO terms were condensed using the *simplify* function (by = "NES", cutoff = 0.7) to minimize redundancy between similar terms. Terms with $P_{adj} < 0.05$ were considered significant and ranked according to their normalized enrichment score (NES, positive indicates upregulation of a pathway, negative indicates downregulation).

*Primary DRG neuron culture and* $Ca^{2+}$ *imaging*

During imaging, cultures were treated with CXCL16 +/- pre-treatment with a CXCR6 antagonist, ML339. For wells receiving CXCL16 only, without ML339, coverslips were stimulated with vehicle, followed by CXCL16 (50 ng/mL), and finally a positive control (KCl, 100 mM). For each stimulation factor, ~750uL was manually removed with a 1000 µl pipette before the stimulation and then that volume was added back with the stimulation factor included. In a similar fashion, wells receiving CXCL16 + ML339 were pre-treated with ML339 (180 µM), followed by addition of stimulation factors – vehicle + ML339, then CXCL16 + ML339, and finally KCl. The initial ML339 bath was left in the chamber throughout the course of the experiment and the concentration of CXCL16 was increased to 100 ng/mL to account for the increased bath volume. Cell viability was assessed at the end of each coverslip experiment with propidium iodide staining in buffer solution (1:3000 dilution).

*Statistical analysis*

For the vehicle vs CXCL16 acute knee hyperalgesia experiment, a two-way linear mixed-effects model compared longitudinally measured, repeated/live, continuous outcomes by sex in the vehicle injected and CXCL16 injected limbs (limb and timepoint as within-subject factors). Knee withdrawal threshold ratio overtime was compared using a one-way linear mixed effects model (timepoint as within-subject factor).

For the CXCL16 vs CXCL16 + ML339 acute knee hyperalgesia experiment, a three-way linear mixed-effects model compared longitudinally-measured, repeated/live, continuous outcomes in the vehicle injected and treatment injected limbs, with treatment being either CXCL16 or CXCL16 + ML339 (limb and timepoint as within-subject factors, and treatment as between-subject factor). Knee withdrawal threshold ratio overtime was compared using a two-way linear mixed effects model (timepoint as within-subject factor, treatment as between-subject factor).

For the acute knee hyperalgesia experiment post-ACLR, a two-way linear mixed-effects model compared longitudinally-measured, repeated/live, continuous outcomes by limb (contralateral or ACLR) in the vehicle treated and ML339 treated mice (timepoint as within-subject factor, treatment as between-subject factor). The opposing limb was factored in as a covariate (i.e. the contralateral limb served as a covariate for the ACLR limb and vice versa). Knee withdrawal threshold ratio overtime was compared using a two-way linear mixed effects model (timepoint as within-subject factor, treatment as between-subject factor). Normality and homoscedasticity of residuals were confirmed for all linear models and all linear models used Sidak post-hoc correction.

### SUPPLEMENTAL FIGURES

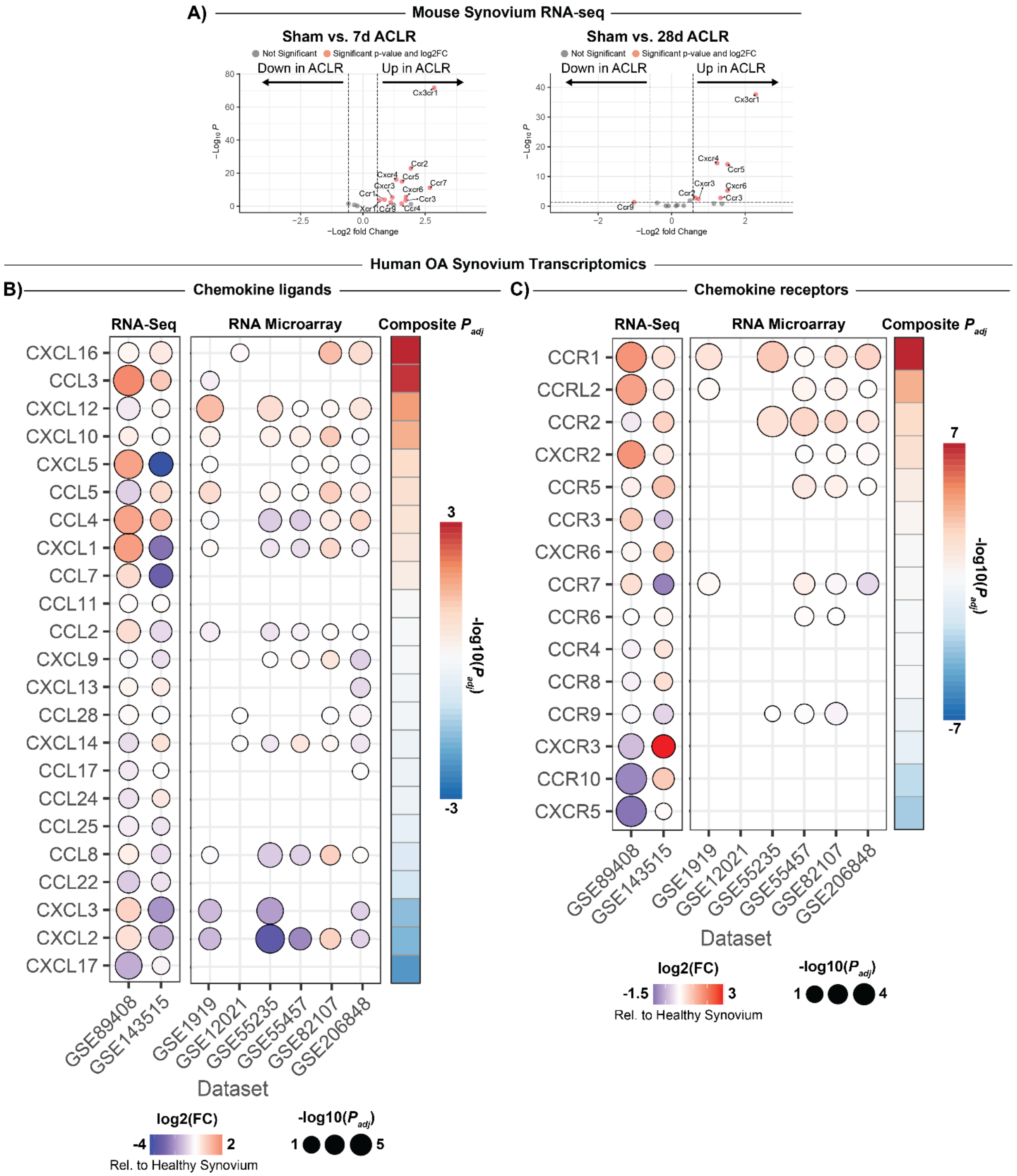


**Fig. S1. Synovial gene expression of chemokine ligands and chemokine receptors is perturbed in mouse and human OA**. (**A**) Volcano plots of mouse synovial bulk RNAseq data *(32)* comparing chemokine receptor expression in Sham vs 7d ACLR and Sham vs 28d ACLR.

Genes with red datapoints were differentially expressed at $P_{adj} < 0.05$ and $|\log_2 FC| > 0.585$. Bubble plots from a meta-analysis of human synovial transcriptomic datasets comparing (**B**) chemokine ligand expression and (**C**) chemokine receptor expression in human OA synovium relative to healthy synovium.

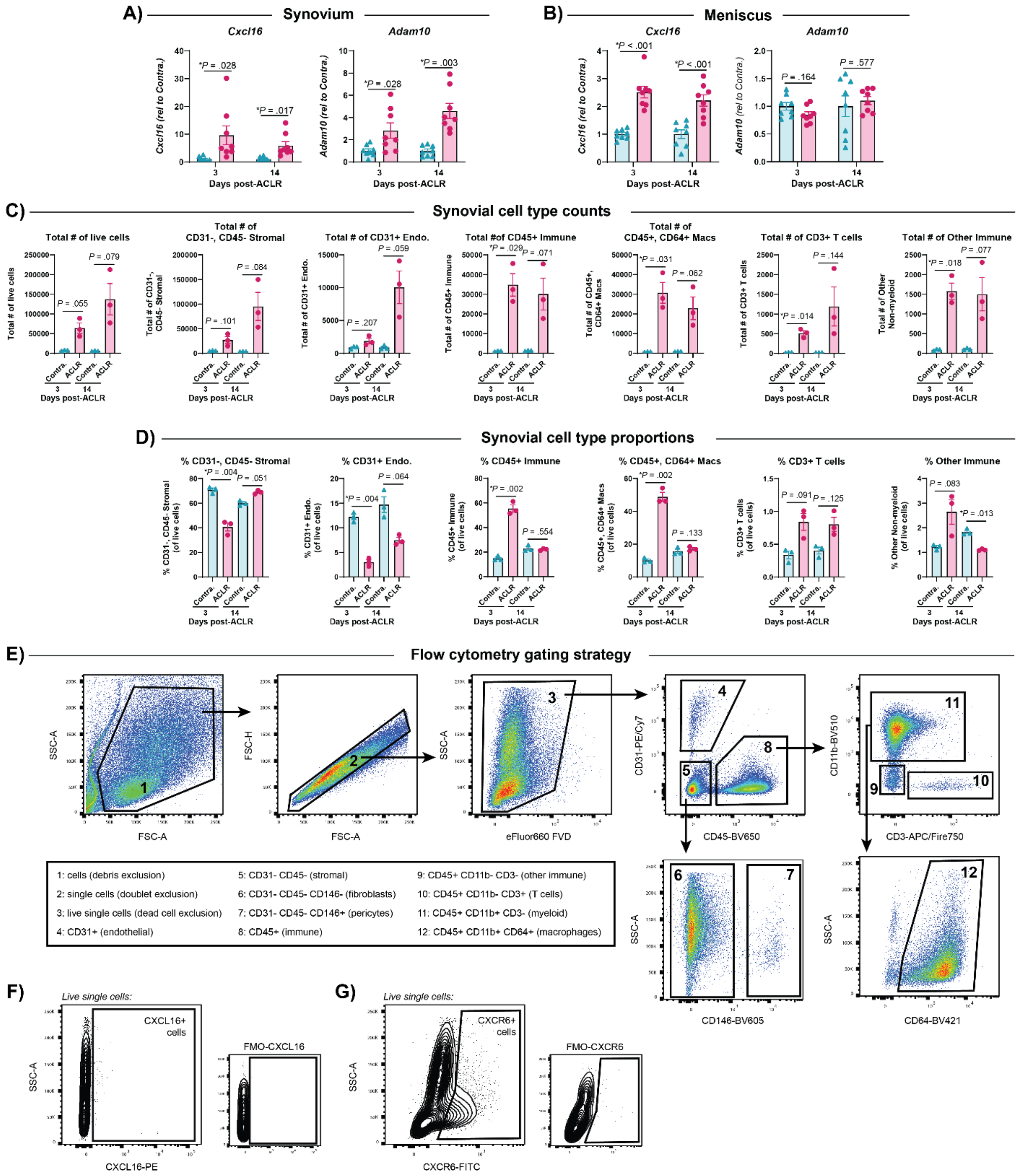


**Fig. S2. qPCR corroboration of CXCL16 signaling axis expression and flow cytometry supplement**. *Cxcl16* and *Adam10* expression in the (**A**) synovium and (**B**) meniscus of injured and contralateral joints at 3d and 14d post-ACLR as measured by qPCR (n=8 per group). Total number (**C**) and proportion (**D**) of synovial cell types at 3d post-ACLR, 14d post-ACLR, and contralateral samples (n=3 per group where 1 sample = 1 male + 1 female synovia pooled). (**E**)

Flow cytometry gating strategy used to identify synovial cell types, with numbers showing the populations in each gate. Flow cytometry plots showing the positive CXCL16+ (**F**) or CXCR6+ (**G**) cells and their corresponding fluorescence-minus-one (FMO) controls. All bars show mean ± SEM.

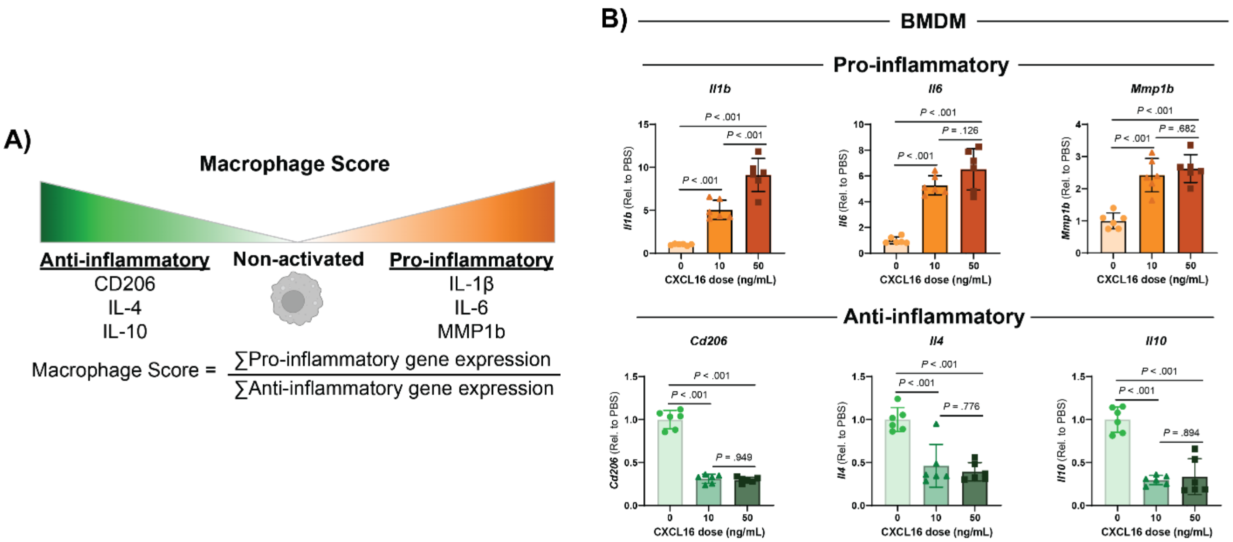


**Fig. S3. CXCL16 treatment *in vitro* – individual expression of macrophage score genes**. (**A**) Macrophage polarization score is calculated by the ratio of pro-inflammatory (*Il1b*, *Il6*, *Mmp1b*) gene expression to anti-inflammatory gene expression (*Cd206*, *Il4*, *Il10*). (**B**) Individual gene expression of each gene within the macrophage score is shown. All bars show mean ± SEM.

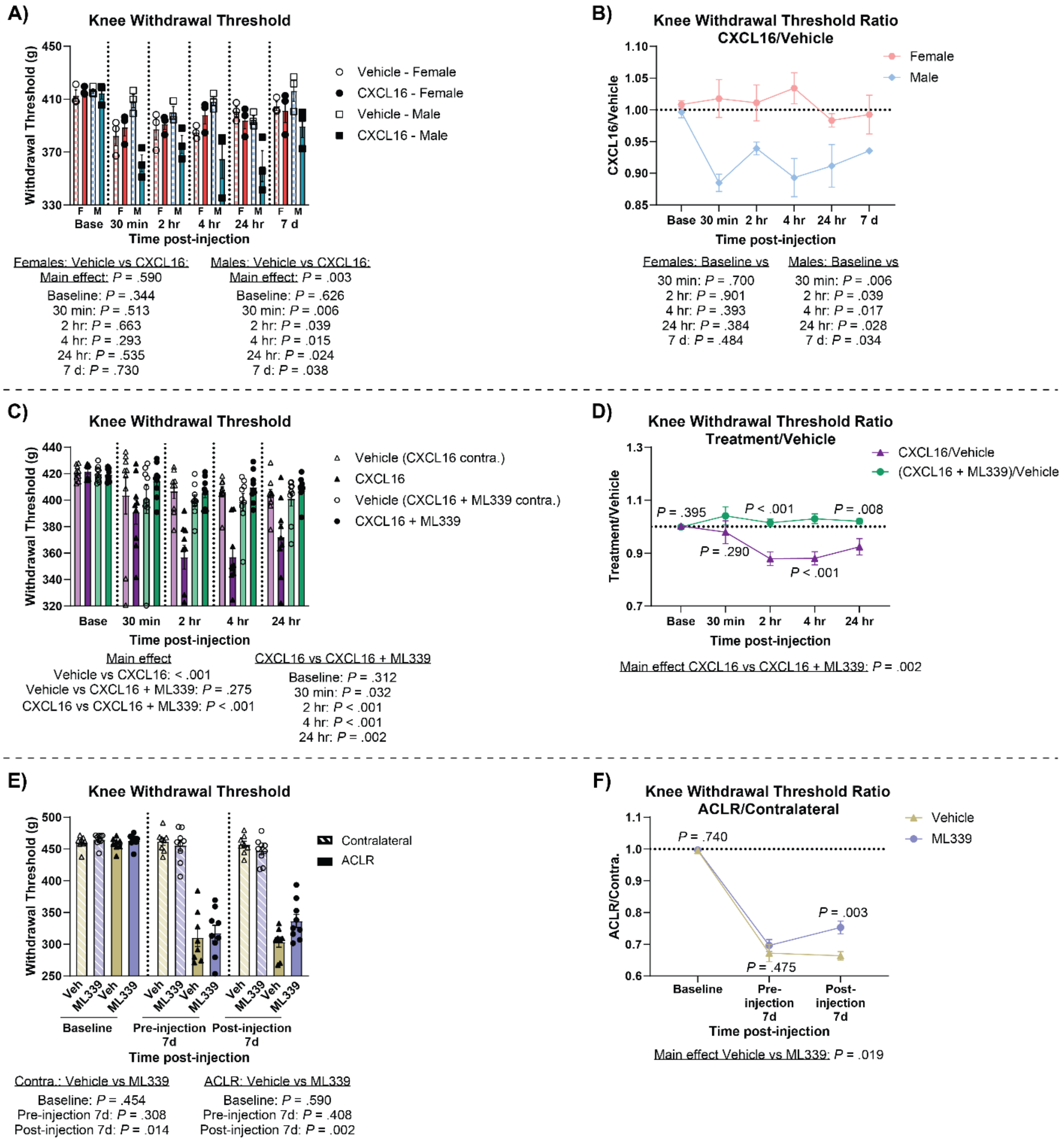


**Fig. S4. Acute knee hyperalgesia supplement.** Knee withdrawal threshold with individual data points shown (**A**) and knee withdrawal threshold ratio summary line plot (**B**) at acute timepoints following intra-articular injection of CXCL16 or vehicle in the contralateral joint (n=3 per sex). Knee withdrawal threshold with individual data points shown (**C**) and knee withdrawal threshold ratio summary line plot (**D**) at acute timepoints following intra-articular injection of CXCL16 or CXCL16 + ML339, and vehicle in the contralateral joint (n=9 per group). Knee withdrawal threshold with individual data points shown (**H**) and knee withdrawal threshold ratio summary line plot (**I**) pre- and post-intraperitoneal injection of vehicle or ML339 at 7d post-ACLR (n=8-9 per group). All bars show mean + SEM.

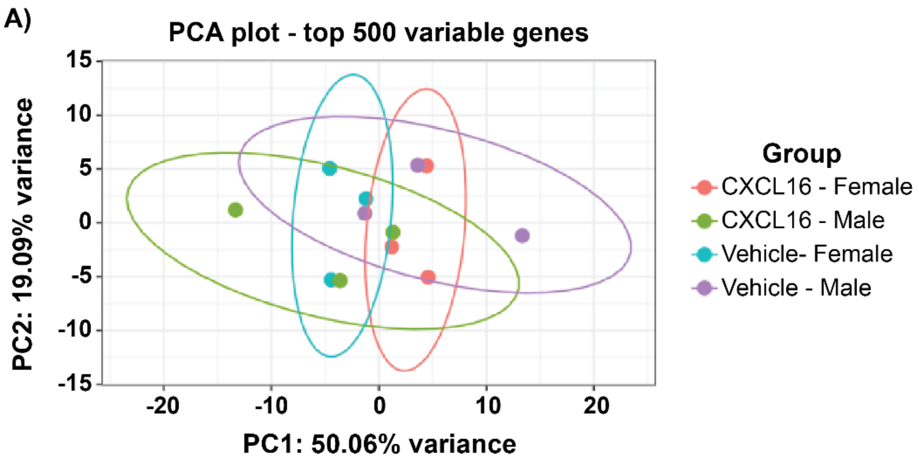


**Fig. S5. Bulk RNAseq PCA plot.** (**A**) Principal component analysis (PCA) plot showing samples grouped by sex and treatment group with 95% confidence ellipses. PCA was computed from VST normalized data (Limma *vst* function), and Limma *removeBatchEffect* function was used to regress out inter-subject variability between pairs of ipsilateral/contralateral limbs.

### SUPPLEMENTAL TABLES

**Table S1. Antibodies.**

| **Antibody** | **Clone (RRID)** | **Vendor** | **Application** | **Dilution** |
|---|---|---|---|---|
| Anti-mouse CD3 AlexaFluor700 | 500A2 (AB_396972) | BD Pharmingen | Flow cytometry | 1:100 |
| Anti-mouse CD3 APC/Fire750 | 17A2 | Biolegend | Flow cytometry | 1:100 |
| Anti-mouse CD11b Brilliant Violet 510 | M1/70 (AB_2561390) | Biolegend | Flow cytometry | 1:200 |
| Anti-mouse CD31 PE/Cy7 | 390 (AB_830756) | Biolegend | Flow cytometry | 1:100 |
| Anti-mouse CD45 Brilliant Violet 650 | 30-F11 (AB_2565884) | Biolegend | Flow cytometry | 1:400 |
| Anti-mouse CD64 Brilliant Violet 421 | X54-5/7.1 (AB_2562694) | Biolegend | Flow cytometry | 1:200 |

| | | | | |
|---|---|---|---|---|
| Anti-mouse CD90/Thy1 PE/Dazzle594 | 30-H12 (AB_2632886) | Biolegend | Flow cytometry | 1:200 |
| Anti-mouse CD146 Brilliant Violet 605 | ME-9F1 (AB_2740163) | BD OptiBuild | Flow cytometry | 1:100 |
| Anti-mouse CXCL16 PE | 12-81 (AB_2869842) | BD Pharmingen | Flow cytometry | 1:100 |
| Anti-mouse CXCR6 FITC | SA051D1 (AB_2572144) | Biolegend | Flow cytometry | 1:100 |

**Table S2. Gene primers for qPCR.**

| ***Gene*** | **Forward primer** | **Reverse primer** |
|---|---|---|
| *Adam10* | GGGCTGGGAGGTCAGTATGG | GTGAGACTGCTCGTTTGGCA |
| *Atp5b* | CTGGATTCAGGGGCACCAAT | GCACCTCCAAAGAGTCCGAT |
| *Cd206* | GGCTGATTACGAGCAGTGGA | ATGCCAGGGTCACCTTTCAG |
| *Cxcl16* | CTTTTCTTGTTGGCGCTGCT | GGACTGCAACTGGAACCTGA |
| *Cxcr6* | GGGCTTCTCTTCTGATGCCA | CTACCAGGTACACACAGGGC |
| *Gapdh* | GCCTCTCTTGCTCAGTGTCC | CTCCCACTCTTCCACCTTCG |
| *IL10* | GCGCTGTCATCGATTTCTCC | ATGGCCTTGTAGACACCTTGG |
| *IL1b* | TGCCACCTTTTGACAGTGATG | AAGGTCCACGGGAAAGACAC |
| *IL1ra* | AGGTGTCTTCTGCTCTACCA | AAGTGACTTGATTGGTCTGG |
| *Il4* | ATTGATGGGTCTCAACCCCC | CTCTGTGGTGTTCTTCGTTGC |
| *Il6* | GCCTTCTTGGGACTGATGCT | TGCCATTGCACAACTCTTTTCT |
| *Mmp1b* | CAGGCCTTATATGGACCTTCC | AGGTCTATCACATCGATCAAAGGT |

### Table S3. PTOA severity scoring.

| PTOA severity scoring | | | | | | | | |
|---|---|---|---|---|---|---|---|---|
| **Category**<br>**Range:**<br>**Region(s):** | **0** | **1** | **2** | **3** | **4** | **5** | **6** | **7** |
| **Structural damage**<br>**Range:** 0-7<br>**Region(s):**<br>1. Femur<br>2. Tibia | Normal cartilage. | Roughened surface with small fibrillations, wavy articular surface. | Fibrillations immediately below superficial layer or some loss of laminal surface. | Horizontal cracks or separations between calcified and non-calcified cartilage. | Mild loss of non-calcified cartilage (<10% surface area). | Moderate loss of non-calcified cartilage (10-50% surface area). | Severe loss of non-calcified cartilage (>50% surface area). | Erosion of cartilage to subchondral bone (any percent surface area). |
| **Proteoglycan loss**<br>**Range:** 0-3<br>**Region(s):**<br>1. Femur<br>2. Tibia | Normal cartilage. | Decreased but not complete loss of Safranin-O staining in non-calcified areas. Saf-O loss does not penetrate completely from superficial to deep cartilage. | Saf-O loss penetrates completely from superficial to deep cartilage. Focal loss of Safranin-O staining in non-calcified area (<30% surface area). | Saf-O loss penetrates completely from superficial to deep cartilage. Diffuse loss of Safranin-O staining in non-calcified cartilage (>30% surface area). | | | | |
| **Chondrocyte hypertrophy**<br>**Range:** 0-1<br>**Region(s):**<br>1. Femur<br>2. Tibia | None. | Enlarged chondrocyte lacunae with lack of Saf-O stain around collapsed cell. | | | | | | |
| **Osteophyte size**<br>**Range:** 0-3<br>**Region(s):**<br>1. Femur<br>2. Tibia | None. | Small – ≤1x thickness as adjacent cartilage. | Medium – >1x to 3x as thick as adjacent cartilage. | Large - >3x thicker than adjacent cartilage. | | | | |
| **Osteophyte maturity**<br>**Range:** 0-3<br>**Region(s):**<br>1. Femur<br>2. Tibia | None. | Predominately cartilage. | Mixed cartilage and bone with vascular invasion. | Predominately bone. | | | | |
| **Subchondral bone thickening**<br>**Range:** 0-3<br>**Region(s):**<br>1. Femur<br>2. Tibia | Normal SCB. | Mild thickening, <50% increase. | Moderate thickening, 50-100% increase. | Severe thickening, >100% increase. | | | | |

### Table S4. Synovitis severity scoring.

| Synovitis severity scoring | | | | |
|---|---|---|---|---|
| **<u>Category</u>**<br>**Range:**<br>**Region(s):** | **0** | **1** | **2** | **3** |
| **<u>Pannus</u>**<br>**Range:** 0-3<br>**Region(s):**<br>1. Anterior synovium/tibia | None. | Mild: Pannus has migrated onto bone but is not encroaching on articular surface. It is in the transition zone where there is calcified cartilage, but you are not yet at the articular surface. | Moderate: Pannus has migrated through the transition zone and <1x cartilage depth onto the articular surface. | Severe: Pannus has migrated > 1x cartilage depth onto the articular surface. |
| **<u>Bone erosion</u>**<br>**Range:** 0-3<br>**Region(s):**<br>1. Anterior femur<br>2. Anterior tibia | None. | Partial thickness loss of cortical bone only. Wavy surface. | Focal complete loss of cortical bone - communication with marrow cavity at one small vascular communication site. A single large “offshoot” of cortical bone erosion. | Widespread complete loss of cortical bone - communication with marrow cavity at multiple sites or broad area loss of cortical bone. |
| **<u>Synovial lining hyperplasia</u>**<br>**Range:** 0-1<br>**Region(s):**<br>1. Anterior, superior synovium<br>2. Anterior, inferior synovium | 1 cell thick. | Mild: 2-3 cells thick. | Moderate: 4-5 cells thick. | Severe ≥6 cells thick. |
| **<u>Subsynovial inflammation</u>**<br>**Range:** 0-3<br>**Region(s):**<br>1. Anterior synovium | None. | One pocket of densely associated inflammatory cells. | Two to three pockets of densely associated inflammatory cells. These are focal areas of dense subsynovial WBC infiltrate – but still predominantly normal subsynovial areolar connective tissue present. | ≥ Four pockets of densely associated inflammatory cells. This is widespread dense subsynovial WBC infiltrate with markedly reduced or little/no normal areolar connective tissue evident and some lymphoid follicle formation. |
| **<u>Synovial fibrosis</u>**<br>**Range:** 0-3<br>**Region(s):**<br>1. Anterior synovium | None (less than 10% to account for the immediate sublining and regular matrix around blood vessels). | Dispersed fibrosis (10% to 1/3 of synovial area). | Moderate fibrosis (>1/3 to 2/3 of synovial area). | Severe fibrosis (>2/3 of synovial area). |
| **<u>Synovial exudate</u>**<br>**Range:** 0-1<br>**Region(s):**<br>1. Anterior synovium | None. | Infiltration of inflammatory cells (neutrophils, macrophages, and/or lymphocytes) or fibrin in the synovial cavity. | | |

### SUPPLEMENTAL DATA FILES

Data file S1. Synovial bulk RNA-seq – quality control.

Data file S2. Synovial bulk RNA-seq – DEG lists.
